# Anti-Parasitics with a Triple Threat: Targeting Parasite Enzymes, the Proton Motive Force, and Host Cell-Mediated Killing

**DOI:** 10.1101/2025.02.03.636145

**Authors:** Akanksha M. Pandey, Satish R. Malwal, Mariana Valladares-Delgado, Liesangerli Labrador-Fagúndez, Bruno G. Stella, Luis José Díaz-Pérez, André Rey-Cibati, Davinder Singh, Marianna Stampolaki, Sangjin Hong, Robert B. Gennis, Antonios Kolocouris, Gustavo Benaim, Eric Oldfield

## Abstract

We investigated the effects of the tuberculosis drug candidate SQ109 (**8a**) and of its analog MeSQ109 (**8b**) against *Leishmania mexicana* in promastigote and amastigote forms, as well as against host cell macrophages finding potent activity (1.7 nM) for MeSQ109 against the intracellular forms, as well as low toxicity (∼61 µM) to host cells, resulting in a selectivity index of ∼36,000. We then investigated the mechanism of action of MeSQ109 finding that it targeted parasite mitochondria, collapsing the proton motive force, as well as targeting acidocalcisomes, rapidly increasing the intracellular Ca^2+^ concentration. Using an *E. coli* inverted membrane vesicle assay, we investigated the pH gradient collapse for SQ109 and 17 analogs finding that there was a significant correlation (on average R=0.67, p∼0.008) between pH gradient collapse and cell growth inhibition in *Trypanosoma brucei, T. cruzi, L. donovani* and *Plasmodium falciparum*. We also investigated pH gradient collapse with other anti-leishmanial agents: azoles, antimonials, benzofurans, amphotericin B and miltefosine. The enhanced activity against intracellular trypanosomatids is seen with *Leishmania* spp. grown in macrophages but not with *Trypanosoma cruzi* in epithelial cells and is proposed to be due in part to host-based killing, based on the recent observation that SQ109 is known to convert macrophages to a pro-inflammatory (M1) phenotype.

---

Leishmaniases are parasitic diseases transmitted by female sandflies that transmit the disease between mammalian hosts and are caused by at least 20 different species of the genus *Leishmania*. Distinct *Leishmania* species induce a spectrum of clinical manifestations, from self-resolving cutaneous lesions to severe visceral disease. The clinical outcome is influenced by the interaction of parasite characteristics, vector biology, and host factors, with immune response being the most critical among the host factors. Cutaneous leishmaniasis typically presents as an ulcerative lesion that undergoes spontaneous resolution within 3 to 18 months. However, it can result in sequelae such as scarring, disfigurement, and social stigmatization. Depending on the *Leishmania* species involved, up to 10% of cutaneous leishmaniasis cases may progress to more severe forms, including mucocutaneous leishmaniasis, diffuse cutaneous leishmaniasis, disseminated cutaneous leishmaniasis, and leishmaniasis recidivans.^1^

Leishmaniasis is currently managed with drugs such as amphotericin B, miltefosine, and the antimonials pentostam and glucantime—which were introduced ∼ 80 years ago. The emergence of drug resistance and the toxicity of current treatments underscore the need for the development of safer, cheap and more efficacious therapies to address *Leishmania* pathogens^2^, in particular drug-resistant strains, and the World Health Organization recommend two newer therapeutics, paromomycin and liposomal amphotericin B (AmBisome), which is however, very expensive. In earlier work, we investigated the effects of the bisphosphonate drug pamidronate, which targets isoprenoid biosynthesis, on mice infected with *Leishmania mexicana amazonensis,* obtaining a radical cure. ^2, 3^ In later work, we investigated the *in vitro* effects of amiodarone, and dronedarone, Figure 1, in *L. mexicana* as well as in *Trypanosoma cruzi*.^4–9^

**Figure 1.**
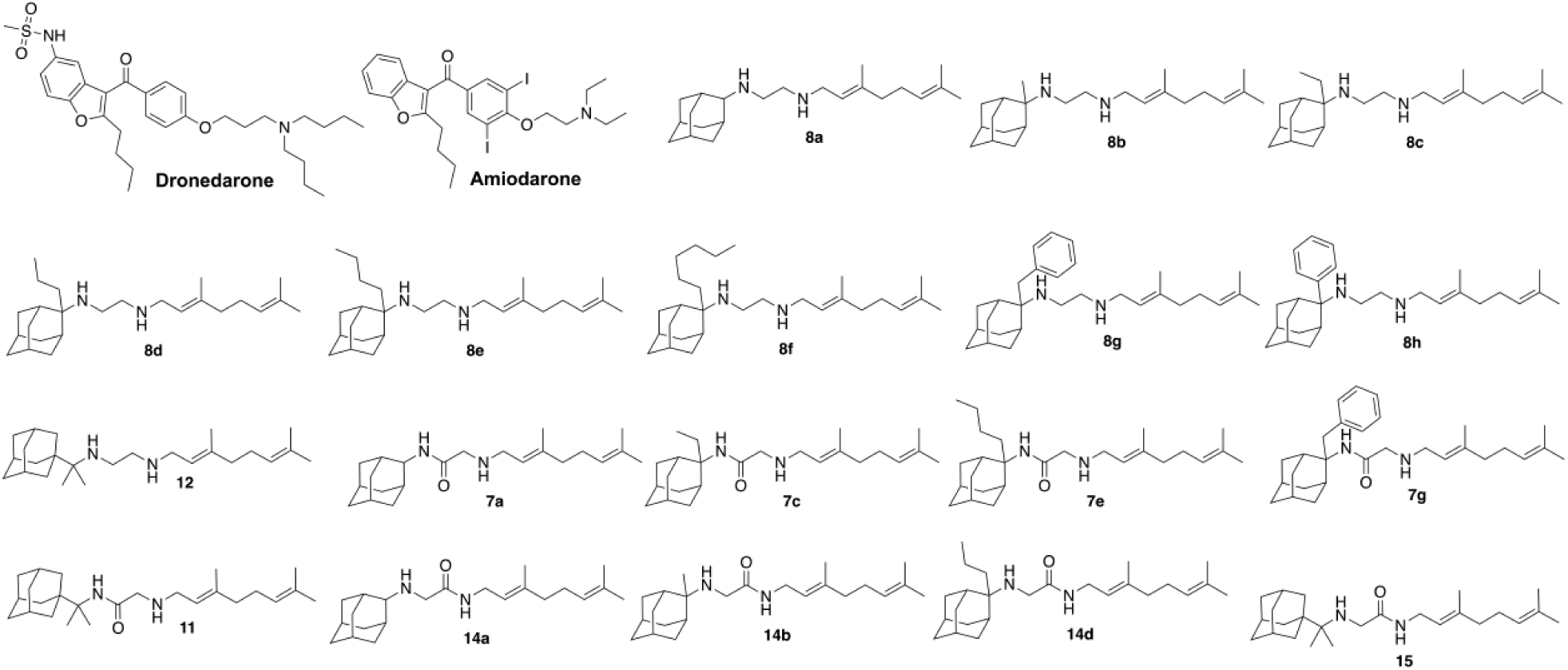
Structures of amiodarone, dronedarone, SQ109 (**8a**), MeSQ109 (**8b**), and 16 other SQ109 analogs investigated in this work. Compound code numbers are those reported in ref. 14.

This work was stimulated by earlier observations that amiodarone had broad spectrum activity against other pathogens—fungi,^10^ that it acted synergistically with azoles that inhibit ergosterol biosynthesis, that it disrupts Ca^2+^ homeostasis,^11^ and the fact that amiodarone was already used to treat patients with Chagas disease, by reducing ventricular arrhythmias. Amiodarone and the related benzophenone dronedarone, are cationic, lipophilic drugs which affect Ca^2+^ homeostasis, the proton motive force as well as sterol biosynthesis, in *Trypanosoma cruzi* as well as *L. mexicana, in vivo.* In other recent work, we investigated the effects of another cationic lipophilic drug, SQ109, a tuberculosis drug candidate, and several analogs, on *T. cruzi* and *L. mexicana* growth *in vitro*,^12, 13,14^ and with SQ109, *in vivo*.^15^

SQ109, *N*^1^-(adamantan-2-yl)-*N*^2^-[(2*E*)-3,7-dimethylocta-2,6-dien-1-yl]ethane-1,2-diamine (**8a**), is an ethylenediamine antitubercular compound that completed phase IIb trials for tuberculosis^16^ and effectively targets multidrug-resistant *Mycobacterium tuberculosis* (Mtb). It targets the mycobacterial membrane protein large 3 (MmpL3), a transporter of trehalose monomycolate (TMM), which is crucial for cell wall biosynthesis. SQ109 has also been implicated in the disruption of the proton motive force (PMF), by functioning as an uncoupler,^17,18^ as well as targeting isoprenoid biosynthesis.^19^ In Trypanosomatid parasites, it has been shown, at least in *T. cruzi*, that SQ109 also has effects on sterol biosynthesis, targeting a side-chain reductase and a Δ^8^, Δ^7^-isomerase,^12^ and it is likely that the corresponding proteins in *L. mexicana* may also be targeted. Additionally, effects on Ca^2+^ homeostasis with SQ109 have been observed, in *T. cruzi*, *L. mexicana* and *L. donovani*, due to targeting of acidocalcisomes and mitochondria. Acidocalcisomes are acidic compartments that store Ca^2+^ and polyphosphates (polyP). They are found in diverse organisms, including trypanosomatids, where they were first characterized.^20^ PolyP binds to both inorganic and organic anions, storing these compounds, which are essential as cofactors for numerous enzymatic reactions, protein synthesis, metabolism, osmoregulation and Ca^2+^ signaling.^21^

Both *Trypanosoma cruzi* (the causative agent of Chagas disease), and *Leishmania* spp. have two life forms in humans: the clinically most relevant intracellular amastigotes, and extracellular forms: promastigotes in *Leishmania spp*. and epimastigotes in *T. cruzi.* In this work, we investigated the activity of MeSQ109, an analog of SQ109, against *L. mexicana* amastigotes and promastigotes, as well as against the host cell, J774 macrophages, as a probe of toxicity. We chose here to study the 2-adamantyl C2′-Me analog of SQ109, MeSQ109, since SQ109 itself is very rapidly metabolized by the liver while MeSQ109 is ∼3x more stable, in liver microsomal assays, and had no effect on HepG2 cells (at a 20 μM level).^14^ We first determined the effects of MeSQ109 on the proton motive force (PMF) (the mitochondrial membrane potential, Δψ, and the pH gradient collapse, ΔpH), and on Ca^2+^ homeostasis, in promastigotes. We then expanded our study to investigate the effects of SQ109 and 17 analogs (Figure 1) on ΔpH using the *E. coli* inverted membrane assay described previously,^17^ correlating these results with cell growth inhibition in a range of protozoan parasites where the activity of these 18 compounds has been reported: ^14^ *T. brucei*, *T. cruzi*, *L. donovani*, as well as against *Plasmodium falciparum*, the malaria parasite. In addition, we investigated the effects of azoles, antimonials, benzofurans, amphotericin B and miltefosine on ΔpH. We then compared our results for intra- and extra-cellular *Leishmania* and *T. cruzi* parasite cell growth inhibition since the former reside inside macrophages and are grown *in vitro* in macrophages, while *T. cruzi in vitro* are typically grown in epithelial cells, either Vero or LLC-MK_2_, and SQ109 is now known to convert macrophages to an M1 phenotype,^22^ similar to the effects seen with miltefosine against *L. donovani* growth in macrophages.^23^

## RESULTS AND DISCUSSION

### Effects of MeSQ109 on *Leishmania mexicana* promastigotes, amastigotes and macrophages

We first characterized the effect of MeSQ109 on the viability of *L. mexicana* promastigotes, using an MTT reduction assay. The results are shown in Figure 2a. To evaluate the effect of MeSQ109 on *L. mexicana* amastigotes internalized in J774 macrophages, infected macrophages were incubated for 72 hours with increasing concentrations of the compound. After this period, the percentage of macrophages containing amastigotes was determined. MeSQ109 had a dose-dependent effect on the number of infected macrophages (Figure 2b). The IC_50_ for *L. mexicana* promastigote cell growth inhibition by MeSQ109 was ∼1.6 μM while that for the amastigotes was ∼1000-fold less (1.7 ± 0.2 nM), even lower than that reported for SQ109 in *L. mexicana* amastigotes where the IC_50_ was 11 ± 0.9 nM.^13^ MeSQ109 also had very little effect on the viability of the macrophages (IC_50_ value ∼ 61 ± 3.9 μM). By way of comparison, the IC_50_ of SQ109 against J774 macrophages (after 72 h of treatment) is much less, 5.8 ± 0.1 μM.^13^ To assess toxicity, we used a selectivity index, SI, given by SI = IC_50_ (host cell)/IC_50_ (parasite cell) finding that for the clinically relevant amastigote form of *L. mexicana* the SI was ∼36,000, a high value. However, SQ109 itself has a SI of 527,^13^ similar to that reported for *L. donovani* amastigotes where SQ109 has a SI of 809.^24^ MeSQ109 has, therefore, potent activity against the parasites as well as low host cell toxicity. Next, we investigated some possible mechanisms of action of MeSQ109, in *L. mexicana*.

**Figure 2.**
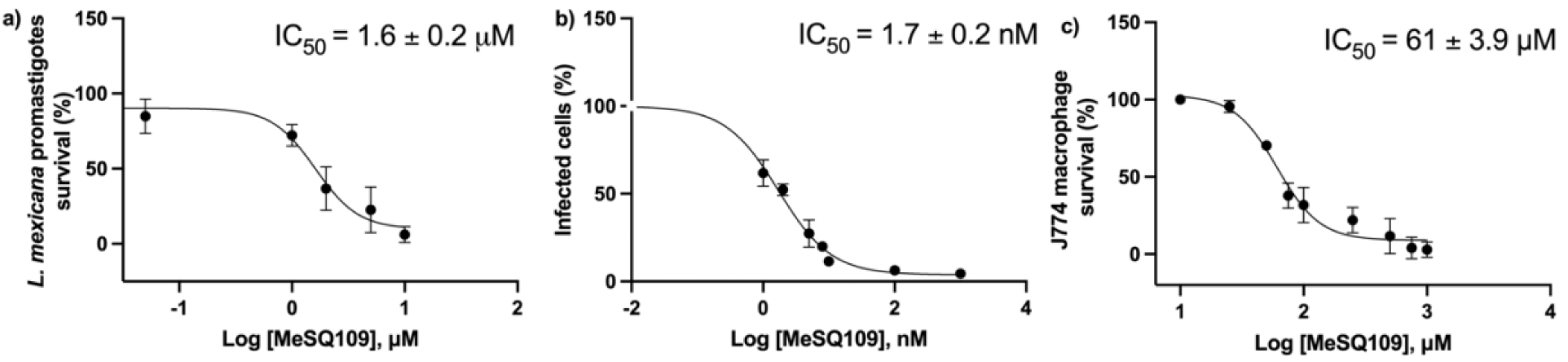
Effects of MeSQ109 on *L. mexicana* promastigote, amastigote and J774 macrophage cell growth inhibition. a) Promastigotes. b) Intracellular amastigotes in *L. mexicana*-infected J774 macrophages. The percentage of infected cells (squares) and the IC_50_ values were determined 72 h post-treatment. c) J774 macrophages. Each bar represents the standard deviation of the experimental points from at least three independent experiments.

### Effect of MeSQ109 on the mitochondrial membrane potential (Δψ) of *L. mexicana* **promastigotes.**

To determine the effect of MeSQ109 on the mitochondrial membrane potential in *L. mexicana* promastigotes, parasites were loaded with the fluorophore rhodamine-123, which accumulates inside mitochondria during respiration, leading to fluorescence quenching. Rhodamine-123 (3,6-diamino-9-[2-(methoxycarbonyl)phenyl]-10λ⁴-xanthen-10-ylium chloride) has a fixed (pyrilium) charge at physiological pH values and its concentration in membranes depends on the membrane potential. Collapse of Δψ induces the release of the fluorophore from the mitochondria and is evidenced by an increase in fluorescence. We show in Figure 3a the result of adding 10 µM MeSQ109 to *L. mexicana* promastigotes loaded with rhodamine-123 in which there is a large increase in fluorescence, indicating collapse of the mitochondrial membrane potential. Further addition of the protonophore FCCP (carbonyl cyanide-p-trifluoromethoxyphenylhydrazone) at 2 µM led to only a minor increase in fluorescence. When FCCP was added before MeSQ109, Figure 3b, there was a large increase in fluorescence, followed by a smaller but reproducible release of the fluorophore upon addition of MeSQ109. The % fluorescence increase on addition of 2, 5 and 10 μM MeSQ109 was then calculated with the maximum increase among the tested concentrations set as 100%, Figure 3c. To make an estimate of the IC_50_ for the collapse of the membrane potential we used the relation:

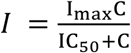

where, I is the % increase in fluorescence, I_max_ is the maximal effect (100 %), C is the concentration and IC_50_ is the concentration for a 50% effect. From the data shown in Figure 3c we find that the IC_50_ is 2.3 ± 0.9 μM. Similar increases in fluorescence for SQ109 itself have been reported^13^ in *L. mexicana* as well as in *L. donovani*^24^. Next, we determined the effects of MeSQ109 on acidocalcisome alkalinization.

**Figure 3.**
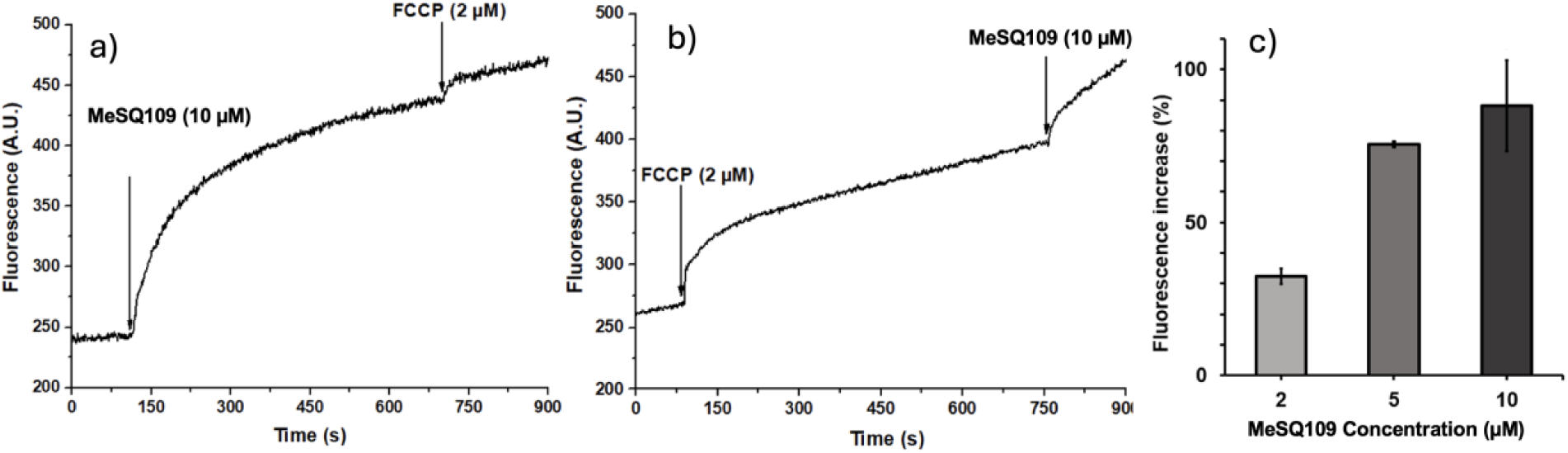
Effect of MeSQ109 on the mitochondrial membrane potential (Δψ_m_) of *L. mexicana* promastigotes. **a)** MeSQ109 (10 μM) was added (arrow) to the parasites previously loaded with rhodamine-123, followed by the addition of FCCP (2 μM). **b)** FCCP (2 μM) was added (arrow) followed by MeSQ109 (10 μM). **c)** The percentage of rhodamine-123 fluorescence increase upon addition of MeSQ109. Bars represent the mean ± SD of three independent experiments. The IC_50_ is 2.3 ± 0.9 μM.

### Effect of MeSQ109 on the alkalinization of acidocalcisomes in *L. mexicana* promastigotes

The effect of MeSQ109 on the alkalinization of acidocalcisomes in *L. mexicana* promastigotes was evaluated by loading the parasites with acridine orange (*N*,*N*,*N*′,*N*′-tetramethylacridine-3,6-diamine). This fluorophore has a pK_a_ of 8.2 and accumulates in acidic compartments, acidocalcisomes, and upon alkalinization it is released, resulting in an increase in fluorescence. Nigericin (2 µM), a K^+^/H^+^ antiporter, was used as a positive control, producing alkalinization of the acidocalcisomes due to the exchange of H^+^ from inside the organelle for cytoplasmic K^+^.

Figure 4a shows the effect of addition of 10 µM MeSQ109 to cells loaded with acridine orange and indicates a large and sustained release of the dye. When nigericin (2 µM) was added after MeSQ109, there was only a very small increase in fluorescence observed, indicating that MeSQ109 induced the almost total alkalinization of the acidocalcisomes. After addition of nigericin (Figure 4b), a large increase in fluorescence was observed. However, it did not reach a maximal response since the addition of MeSQ109 further increased the fluorescence. This may be due to acridine orange accumulating in other acidic inclusions, such as vacuoles or phagolysosomes.^5^ To determine the dose-response effect of MeSQ109 on the degree of alkalinization of *L. mexicana* promastigote acidocalcisomes, the percentage increase in fluorescence was calculated, taking the maximum increase obtained among the tested concentrations as 100%. As can be seen in Figure 4c, there are large increases in fluorescence on increasing the MeSQ109 concentration, resulting from the alkalinization of the acidocalcisomes, and we obtained an IC_50_ of 2.4 ± 0.7 µM for this effect, essentially the same as that observed with the collapse in the mitochondrial membrane potential.

**Fig. 4.**
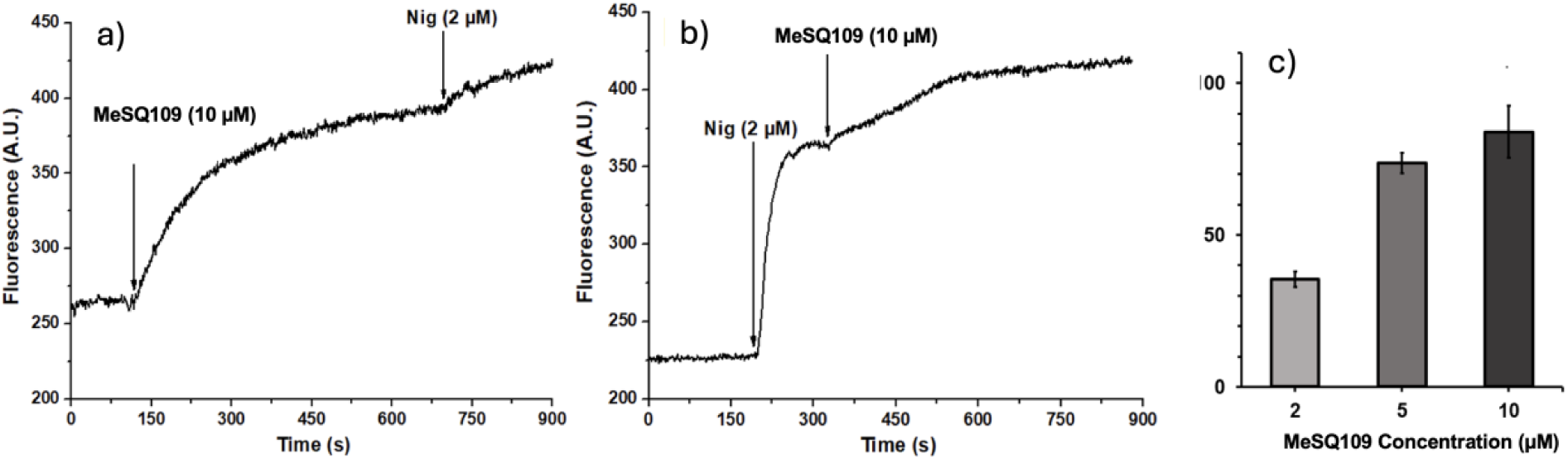
Effects of MeSQ109 on the level of acidocalcisome alkalinization in *L. mexicana* promastigotes. **a)** MeSQ109 at 10 μM was added (left arrow) to parasites loaded with acridine orange, followed by the addition of nigericin at 2 μM (right arrow). **b)** Addition of nigericin (2 μM) was followed by MeSQ109 (10 μM). **c)** The percentage of acridine orange fluorescence increase after the addition of 2, 5 and 10 µM MeSQ109 to *L. mexicana* promastigotes. IC_50_ = 2.4 ± 0.7 µM. Bars represent the mean ± SD of at least three independent experiments.

### Effect of MeSQ109 on the intracellular Ca^2+^ concentration of *L. mexicana* promastigotes

It is known that Ca^2+^ is an essential signaling messenger in all eukaryotic cells, including trypanosomatids, and that homeostasis of the intracellular Ca^2+^ concentration ([Ca^2+^]_i_) is regulated by diverse mechanisms—at the plasma membrane level as well as by intracellular organelles, with high [Ca^2+^]_i_ correlating with cytotoxicity.^25^ In order to determine the effect of MeSQ109 on intracellular Ca^2+^ concentration changes in *L. mexicana* promastigotes, parasites were first loaded with Fura-2 AM, a cell permeable acetoxymethyl analog of the Ca^2+^ indicator Fura-2, an aminopolycarboxylic acid which fluoresces when bound to Ca^2+^ and which forms on hydrolysis of Fura-2 AM by cell esterases. After the addition of 10 μM MeSQ109 (Figure S1), the fluorescence signal at 510 nm increases, indicating that MeSQ109 increases the intracellular Ca^2+^ concentration. Similar results were found for SQ109 itself in *L. mexicana* and *L. donovani* promastigotes.^13, 24^ Digitonin, which permeabilizes the plasma membrane of the parasite, induced a rapid elevation of the intracellular Ca^2+^ concentration, which fell to the minimal value upon the addition of the Ca^2+^ chelating agent, EGTA (Figure S1).

The results discussed above show that the adamantyl-C2′-methyl analog of SQ109, MeSQ109, has a potent effect on the growth of both *L. mexicana* promastigotes, as well as of *L. mexicana* amastigotes, resident inside J774 macrophages, and in previous work, we found that SQ109^13^ and two other cationic lipophilic drugs, amiodarone and dronedarone,^6^ had a similar increase in activity against *L. mexicana*, inside J774 cells. We also showed that amiodarone and dronedarone had major effects on the PMF in promastigotes, acting as uncouplers, as well as on Ca^2+^ homeostasis, similar to the results we found in *L. mexicana*. We also reported that amiodarone, as well as SQ109, had activity against *T. cruzi*, in trypomastigotes, epimastigotes and amastigotes.^12^ In other work, in bacteria, we found that SQ109 had potent activity against mycobacteria, and that there was a correlation between the activity of SQ109 in mycobacteria and its effects on the PMF, affecting both the pH gradient (ΔpH) and the membrane potential (Δψ),^17^as seen here. We also showed that SQ109 and a series of analogs^14^ had potent activity against the malaria parasite, *Plasmodium falciparum*,^26^ and that there was a good correlation between activity (logIC_50_) and logP, the logarithm of the computed oil/water partition coefficient.^14^ There was, however, no correlation with computed logD_7.4_ values.^14^ Taken together, these earlier results suggested that there might be a general correlation between uncoupling activity—collapse of the proton motive force—with activity of SQ109 and its analogs against other pathogenic protozoa: *T. brucei*, *T. cruzi* (epimastigotes), *T. cruzi* (amastigotes), *L. donovani*, as well as against *P. falciparum*. We therefore next determined the effects of SQ109 together with 17 SQ109 analogs (Figure 1) on the pH gradient (ΔpH), using the *E. coli* inverted membrane vesicle (IMV) system with ACMA (9-amino-6-chloro-2-methoxyacridine) fluorescence quenching, as reported previously.^17^ In this earlier work we found that the pH gradient collapse with a series of (other) cationic, lipophilic, *M. tuberculosis* cell growth inhibitors was highly correlated, in the IMV assay, with the Δψ collapse (using an oxonol VI probe) with an R value of ∼ 0.8. Additionally, there was a correlation between ΔpH and Δψ collapse and cell growth inhibition with an R=0.6-0.7.^16, 27^

### SQ109 and its analogs target the PMF in both *Trypanosomatid* and *Apicomplexan* parasites

We determined the effects of SQ109 and the 17 analogs shown in Figure 1 on ΔpH collapse in inverted membrane vesicles, in addition to results for azoles, amphotericin B, antimonials and miltefosine, drugs that are used to treat trypanosomatid infections. Results for SQ109, MeSQ109, amiodarone and dronedarone are shown in Figure 5a-d. As can be seen in Figure 5a and 5b, SQ109 and MeSQ109 had similar activity in collapsing the ΔpH gradient with an IC_50_ of ∼ 1 μM, the same as that found with amiodarone, Figure 5c. Dronedarone was slightly less active (IC_50_ ∼ 2 μM), Figure 5d. Each of these 4 compounds is a lipophilic base and sites of protonation are shown in blue in Figure 5a-d. The known protonophore uncoupler FCCP was about 10 times more active with an IC_50_ of 0.17 μM (data not shown). The collapse of the pH gradient will result in decreased ATP production and is expected to contribute to cell growth inhibition. Next, we investigated whether other drug classes that have anti-parasitic activity had any effects on the proton motive force.

**Figure 5.**
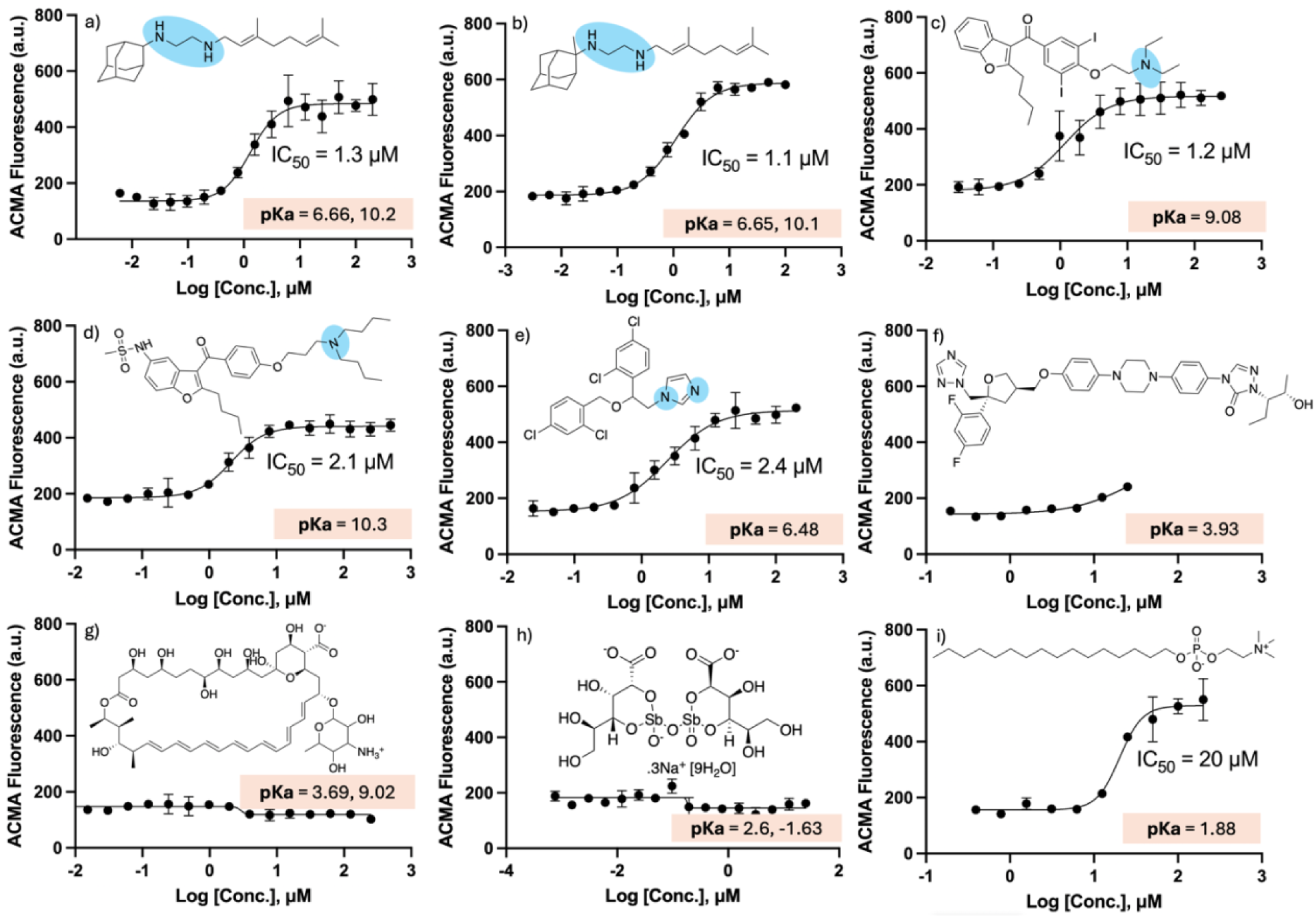
Dose-response curves for pH gradient collapse using *E. coli* IMVs and ACMA fluorescence. a) SQ109. b) MeSQ109. c) amiodarone. d) dronedarone. e) miconazole. f) posaconazole. g) amphotericin B. h) pentostam. i) miltefosine. The blue ellipses show sites that are protonatable in the physiological pH range. Amphotericin and miltefosine are zwitterionic. The salmon-shaded rectangle contains the computed pKa values (chemicalize.com).

In very early work it was shown that azole drugs used to treat fungal infections also had activity against *L. tropica*^28^ and *T. cruzi.*^29^ These early compounds were imidazoles and might be expected to have effects on ΔpH gradient collapse and as can be seen in Figure 5e, this is the case with miconazole which has an IC_50_=2.4 μM, the same as dronedarone. However, there is essentially no effect on the pH gradient with the modern azole drug, posaconazole, Figure 5f. The reason for the difference is that the early azole drugs were imidazoles which have pKa values of ∼6.7 (so can transport protons in a physiological pH range) while the more modern azoles are triazoles and are neutral species. The drug amphotericin (the active component of AmBisome) is also a neutral (zwitterionic) species with computed pK_a_ values of 3.7 (strongest acidic) and 9.0 (strongest basic) so it has no effect on ΔpH, Figure 5g, and as expected, pentostam also has no effect, Figure 5h. Surprisingly, miltefosine, *n*-hexadecyl phosphocholine, is also a neutral, zwitterionic species and cannot directly transport H^+^, but as can be seen in Figure 5i, it does collapse the pH gradient with an IC_50_=20 μM. While this is a high value, the miltefosine concentration in treated *L. donovani* promastigotes has been reported to be ∼1 mM.^30^ Miltefosine is known to reduce ATP formation in *L. donovani*, to decrease the mitochondrial membrane potential and to inhibit cytochrome c oxidase: an effect on ΔpH is expected to contribute to its activity.

Next, we correlated the log IMV (μM) results for SQ109 and the 17 SQ109 analogs (Table 1) with the previously reported logIC_50_ (μM) results for *T. brucei*, *T. cruzi* (epimastigote), *T. cruzi* (amastigote), *L. donovani*,^14^ as well as for *P. falciparum*.^14, 26^ Representative results are shown in Figure 6 and as can be seen, there is significant correlation between the logIC_50_ for cell growth inhibition values and the experimental logIMV results.

**Table 1.**
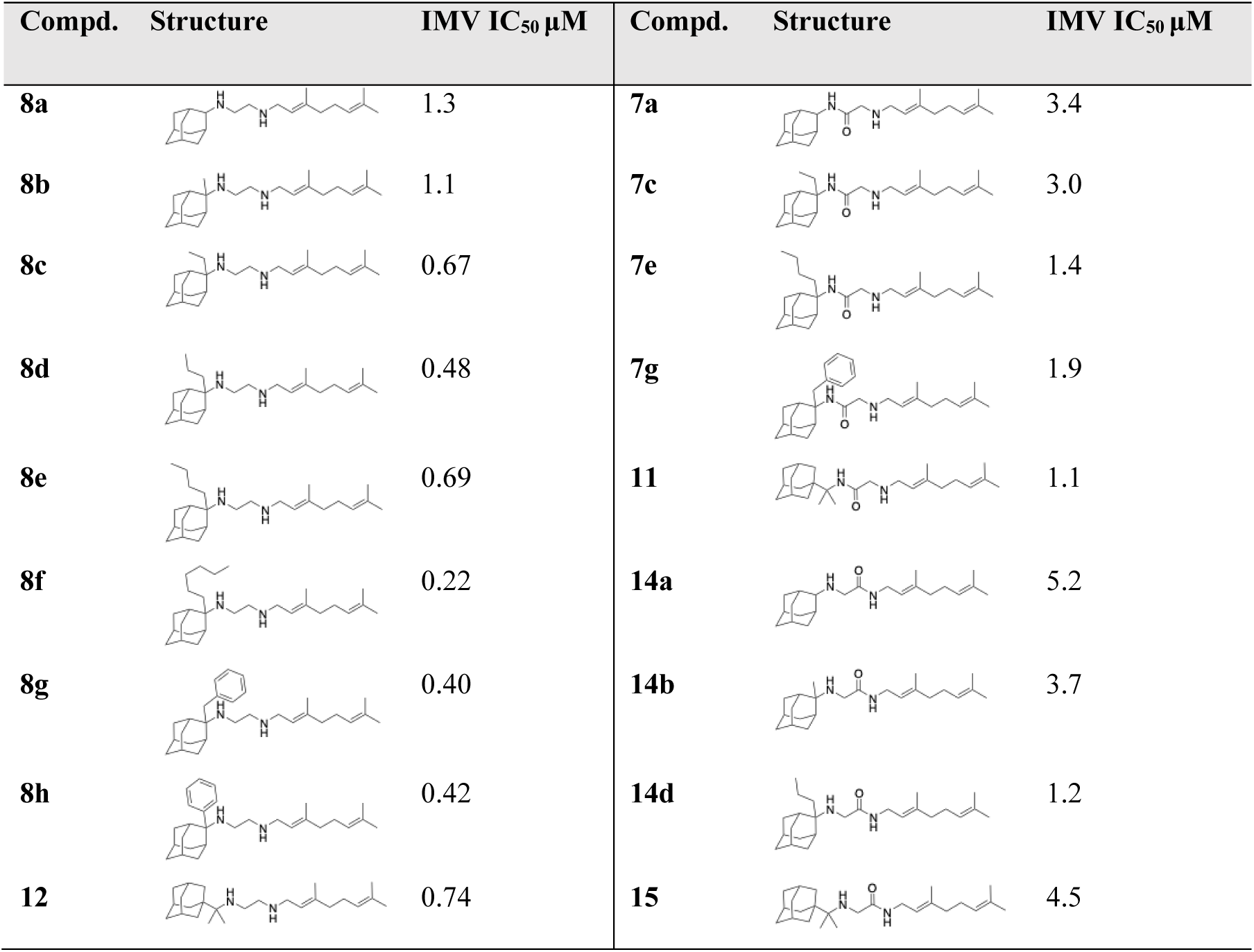
IMV data for SQ109 and its 17 analogs

| Compd. | Structure | IMV IC<sub>50</sub> μM | Compd. | Structure | IMV IC<sub>50</sub> μM |
| --- | --- | --- | --- | --- | --- |
| **8a** |  | 1.3 | **7a** |  | 3.4 |
| **8b** |  | 1.1 | **7c** |  | 3.0 |
| **8c** |  | 0.67 | **7e** |  | 1.4 |
| **8d** |  | 0.48 | **7g** |  | 1.9 |
| **8e** |  | 0.69 | **11** |  | 1.1 |
| **8f** |  | 0.22 | **14a** |  | 5.2 |
| **8g** |  | 0.40 | **14b** |  | 3.7 |
| **8h** |  | 0.42 | **14d** |  | 1.2 |
| **12** |  | 0.74 | **15** |  | 4.5 |

**Figure 6.**
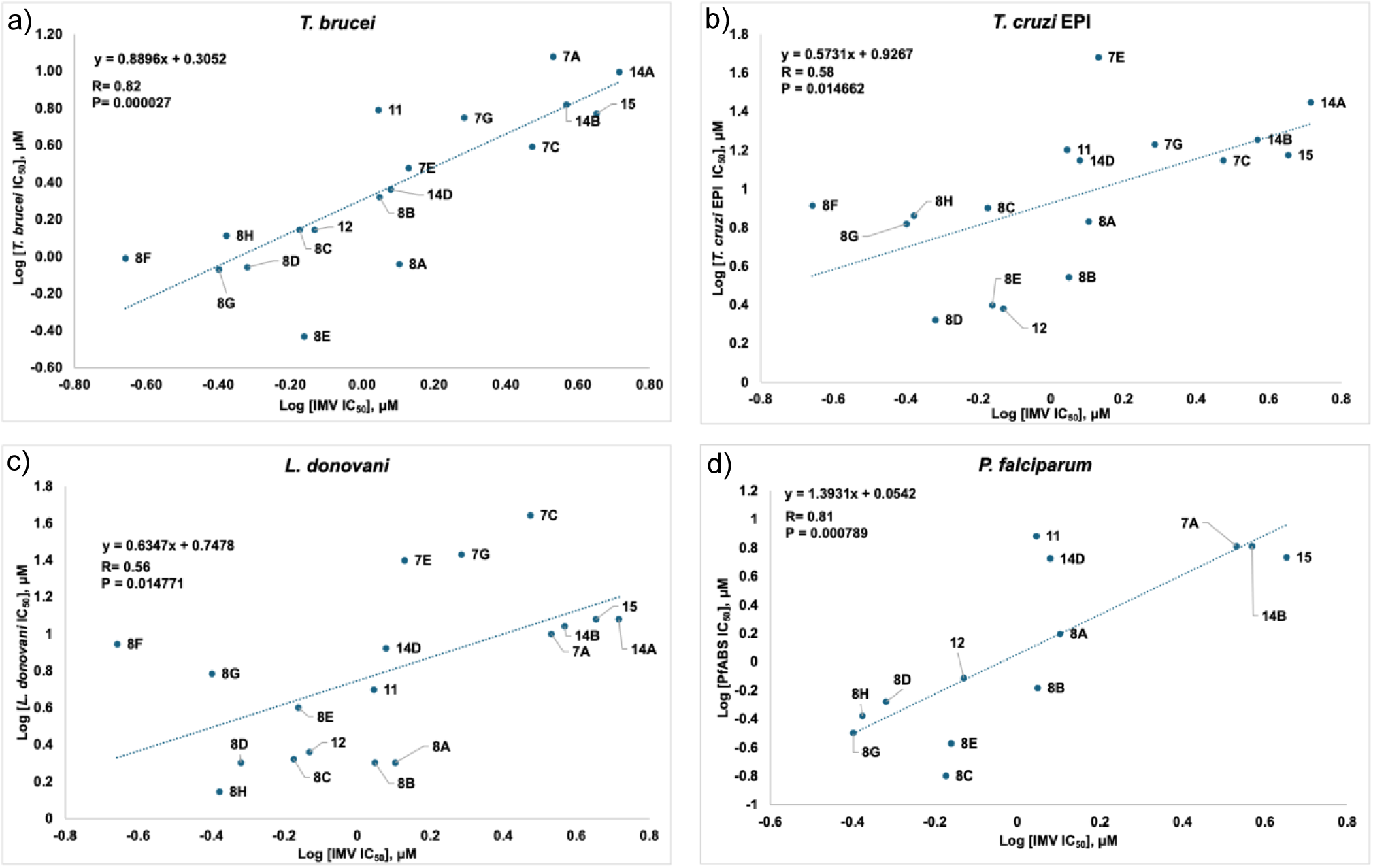
Representative correlation between IMV logIC_50_ and cell growth inhibition values for SQ109 and its 17 analogs. a) *T. brucei.* b) *T. cruzi* epimastigotes. c)*, L. donovani.* d) *Plasmodium falciparum*.

For the correlation between the cell growth inhibition in the 5 protozoa and the IMV pH gradient collapse results the Pearson R-value was on average ∼0.67 and the p-value was ∼ 0.008 (Supporting Information Table S1) over, on average, a ∼ 1.6 x logIC_50_ (μM) range, corresponding to a ∼ 35 x range in activity. Correlation between activity and computed logP values was generally less (Figure S2) and there was no significant correlation with computed logD_7.4_ (Figure S3). This is somewhat surprising since we expected improved results with logD_7.4_, which should give an improved oil/water partition coefficient, but that is not the case. One possibility is that logD_7.4_ calculations for the dibasic SQ109 species are not that accurate, because there are two ionizable groups in close proximity, as well as of course the presence of several charged species around physiological pH values. The results are also of interest since the best correlation was always found between the experimental activity results and the experimental measurements of the pH gradient collapse, as opposed to correlation with a computed property, logP, which could be due, for example, to binding to proteins. Moreover, of course, we do see effects on the pH gradient collapse in *L. mexicana*, as well as in *T. cruzi*, with SQ109, amiodarone as well as with dronedarone. They are also of interest since they strongly suggested the possibility that the effects of SQ109 in the malaria parasite *P. falciparum*, where we previously reported^26^ a correlation between activity and a computed logP, are likely to arise, at least in part, from pH-gradient collapse.

### Activity of SQ109 and analogs against mycobacteria and protozoa

In order to see to what extent the ΔpH collapse effect might contribute to the activity of SQ109 and its analogs against eukaryotes *versus* mycobacteria, the original targets for SQ109 (and its analogs), we used the published cell growth inhibition results^14, 26^ for *T. brucei*, *P. falciparum,* U2OS (a human osteocarcinoma cell line), *T. cruzi* (amastigotes), *T. cruzi* (epimastigotes), *L. donovani*, as well as against three *M. tuberculosis* strains (H37Rv, Erdman and HN878) and *M. smegmatis,* and determined the Pearson correlation coefficients (R) and the probability scores (p-values) between each of the cell growth inhibition results, and with the IMV ΔpH collapse results. As can be seen in Table 2, there are strong correlations (R) between all the eukaryotic cells, on average R=0.78 and p=0.0023, and between the different mycobacteria (on average R=0.85 and p=0.00019) while for the eukaryote/bacteria comparisons the correlations are less: on average R=0.56 and p=0.038. The corresponding matrix of p-values is given in Table S2. These results suggest that effects of ΔpH collapse by SQ109 and its analogs on the eukaryotes are more important than those seen in the mycobacteria, due most likely to the known importance of targeting the MmpL3 protein in the mycobacteria. Of course, while correlation does not imply causation, as we showed above and elsewhere, SQ109, MeSQ109 and other SQ109 analogs do have major effects on the PMF in eukaryotes, as well as in mycobacteria.^17^ Moreover, the effects on the PMF are seen not only with fluorescent probes, but also by using ^31^P NMR chemical shifts to directly determine the pH gradient^17^.

**Table 2.** Correlation between R values for eukaryotes and bacterial cell growth inhibition logIC_50_ (μM) results by SQ109 and a series of 17 analogs, and the results for ΔpH collapse (logIC_50_ IMV)

|  | Eukaryotes |  |  |  |  |  |  | Mycobacteria |  |  |  |
| --- | --- | --- | --- | --- | --- | --- | --- | --- | --- | --- | --- |
|  | IMV | Tb | Pf | U2OS | Tc (A) | Tc (E) | Ld | Mtb H37Rv | Mtb Erd | Ms | Mtb HN878 (IC<sub>50</sub>) |
| IMV | 1 | 0.82 | 0.81 | 0.68 | 0.6 | 0.58 | 0.56 | 0.49 | 0.45 | 0.4 | 0.37 |
| *T. brucei* | 0.82 | 1 | 0.83 | 0.8 | 0.75 | 0.76 | 0.58 | 0.66 | 0.65 | 0.65 | 0.61 |
| *P. falciparum* | 0.81 | 0.83 | 1 | 0.86 | 0.74 | 0.7 | 0.71 | 0.6 | 0.59 | 0.78 | 0.52 |
| U2OS | 0.68 | 0.71 | 0.86 | 1 | 0.9 | 0.95 | 0.72 | 0.58 | 0.58 | 0.6 | 0.55 |
| *T. cruzi* (A) | 0.6 | 0.75 | 0.74 | 0.9 | 1 | 0.9 | 0.8 | 0.58 | 0.58 | 0.66 | 0.75 |
| *T. cruzi* (E) | 0.58 | 0.76 | 0.7 | 0.95 | 0.9 | 1 | 0.81 | 0.49 | 0.46 | 0.38 | 0.57 |
| *L. donovani* | 0.56 | 0.58 | 0.71 | 0.72 | 0.8 | 0.81 | 1 | 0.49 | 0.48 | 0.42 | 0.69 |
| *Mtb* H37Rv | 0.49 | 0.66 | 0.6 | 0.58 | 0.58 | 0.49 | 0.49 | 1 | 0.99 | 0.72 | 0.89 |
| *Mtb* Erd | 0.45 | 0.65 | 0.59 | 0.58 | 0.58 | 0.46 | 0.48 | 0.99 | 1 | 0.75 | 0.93 |
| *Ms* | 0.4 | 0.65 | 0.78 | 0.6 | 0.66 | 0.38 | 0.42 | 0.72 | 0.75 | 1 | 0.83 |
| *Mtb* HN878 (IC<sub>50</sub>) | 0.37 | 0.61 | 0.52 | 0.55 | 0.75 | 0.57 | 0.69 | 0.89 | 0.93 | 0.83 | 1 |

As to possible protein targets for SQ109 and its analogs: in *M. tuberculosis*, a known target for SQ109 is MmpL3, a transporter of TMM. However, this protein is absent in *T. brucei*, *L. donovani*, *L. mexicana*, and *P. falciparum*, although a protein of unknown function having a (predicted) similar 3D and active site structure to the MtMmpL3 is present in *T. cruzi*.^31^ Moreover, TMM is not present in the parasites. Mycobacterial cell growth inhibition by SQ109 is rescued by addition of undecaprenyl phosphate, consistent with inhibition of peptidoglycan biosynthesis, and SQ109 inhibits undecaprenyl diphosphate synthase, albeit weakly,^19^ suggesting a possible role for inhibition of the related isoprenoid biosynthesis in the pathogenic protozoa, involving dolichols. More concretely, SQ109 and amiodarone have been shown to inhibit ergosterol biosynthesis in *T. cruzi*, where they affect an NADPH reductase^12^ and oxidosqualene cyclase,^4^ and dronedarone inhibits ergosterol biosynthesis in *L. mexicana*, also by targeting oxidosqualene cyclase.^5^ However, the effects of SQ109 on the proton motive force are very rapid, a few minutes, while effects on sterol biosynthesis involve multiple enzyme turnovers and are expected to be much slower, on the order of many hours, although they may be significant when cells are grown for days.

### A general proposal for the mechanism of action of SQ109 and some other antiparasitics

What is also of interest about the results presented here and elsewhere is that SQ109, MeSQ109, amiodarone and dronedarone are more effective against intracellular amastigotes than the extracellular forms. For ease of discussion, we show in Table 3 the activity of amiodarone, dronedarone and SQ109 against *T. cruzi* and *L. mexicana,* the activity of MeSQ109 against *L. mexicana*, and of miltefosine, used extensively to treat *L. donovani* infections, against *L. donovani* as well as *L. mexicana*.^32^

**Table 3.** Summary of activities (IC_50_ in μM) of amiodarone, dronedarone, SQ109, MeSQ109 in *T. cruzi*, and *L. mexicana* and host cells, and selectivity index (SI) values

| Cell system | Amiodarone | Dronedarone | SQ109 | MeSQ109 | Miltefosine |
| --- | --- | --- | --- | --- | --- |
| *Tc* EPI IC<sub>50</sub> μM | 9<sup>a</sup> | 4.6<sup>b</sup> | 4.6<sup>c</sup> | - | - |
| *Tc* AMA IC<sub>50</sub> μM | 2.7<sup>a</sup> | 0.75<sup>b</sup> | 0.5-1<sup>c</sup> | - | - |
| Host cell IC<sub>50</sub> μM | NM<sup>a</sup> | NM<sup>b</sup> | 9<sup>c</sup> | - | - |
| *Tc* EPI/*Tc* AMA | 3.3<sup>a</sup> | 6.1<sup>b</sup> | ~8.3<sup>c</sup> | - | - |
| SI | NM<sup>a</sup> | NM<sup>b</sup> | ~10-20<sup>c</sup> | - | - |
| *Lm* PRO IC<sub>50</sub> μM | 0.9<sup>d</sup> | 0.115<sup>e</sup> | 0.53<sup>f</sup> | 1.6 | 7.5<sup>g</sup> |
| *Lm* AMA IC<sub>50</sub> μM | 0.008<sup>d</sup> | 0.00065<sup>e</sup> | 0.011<sup>f</sup> | 0.0017 | 8.5<sup>g</sup> |
| Host cell IC<sub>50</sub> μM | NM<sup>d</sup> | NM<sup>e</sup> | 5.8<sup>f</sup> | **61** | - |
| *Lm* PRO/*Lm* AMA | 113<sup>d</sup> | 177<sup>e</sup> | 48<sup>f</sup> | **941** | 0.88<sup>g</sup> |
| SI | NM<sup>d</sup> | NM<sup>e</sup> | ~500<sup>f</sup> | **~36000** | - |
| *Ld* PRO IC<sub>50</sub> μM | - | - | 0.63<sup>h</sup> | - | 0.41<sup>g</sup> |
| *Ld* AMA IC<sub>50</sub> μM | - | - | 0.0072<sup>h</sup> | - | 3.9<sup>g</sup> |
| Host cell IC<sub>50</sub> μM | - | - | 5.8<sup>h</sup> | - | - |
| *Ld* PRO/*Ld* AMA | - | - | 87.5<sup>h</sup> | - | 0.11<sup>g</sup> |
| SI | - | - | - | - | - |

<sup>a</sup> see ref. 4; <sup>b</sup> see ref. 7; <sup>c</sup> see ref. 12; <sup>d</sup> see ref. 9; <sup>e</sup> see ref. 5, <sup>f</sup> see ref. 13, <sup>g</sup> see ref. 32, <sup>h</sup> see ref. 24. NM=not measured. AMA=amastigote. PRO=promastigote.

With *T. cruzi*, activity against epimastigotes is on average ∼ 6 μM but there is more activity against amastigotes, although the effect is not very large, a ∼3-10 fold increase (Table 3). With (extracellular) *L. mexicana* promastigotes, there is greater activity than that found against (extracellular) *T. cruzi* epimastigotes and substantially more activity with the intracellular *L. mexicana* amastigotes than seen with *T. cruzi* amastigotes. The ratio of promastigote to amastigote activity for amiodarone, dronedarone and SQ109 in *L. mexicana* is on average about 100x, and for MeSQ109 the ratio is even larger, ∼ 900x (Table 3). Similar results were found for SQ109 against *L. donovani*, the causative agent of visceral leishmaniasis, where the IC_50_ against promastigotes is 630 nM while that against amastigotes is much lower, 7.2 nM.^24^ However, MeSQ109 has not only better activity than SQ109 against *L. mexicana* amastigotes but also less host cell toxicity, resulting in a SI ∼ 36,000, as noted above. What, then, might be the differences in intra/extra-cellular activity be due to?

One clear difference between the *T. cruzi* and *L. mexicana* amastigote experiments is that the host cells used in the *L. mexicana* experiments were murine-derived macrophages, J774 cells. In contrast, the results reported previously with *T. cruzi* were obtained with monkey kidney epithelial cells—either Vero cells or LLC-MK_2_—not macrophages. This suggests the possibility that the differences in activity against intracellular/extracellular forms might be due to the host cell as opposed to differences in directly targeting the parasites. Of interest here is the observation that in recent work^22^ with SQ109 inhibition of *M. tuberculosis*, it has been shown that SQ109 targets murine macrophages, converting cells to an M1 (pro-inflammatory) phenotype, resulting in activation of the mitogen-activated protein kinase (MAPK) pathway, a large (∼25x) increase in iNOS production, and downregulation of the M2-specific marker, arginase^22^. The same effects are seen with infected peritoneal macrophages treated with SQ109, and SQ109-pretreated macrophages effectively kill *M. tuberculosis*, although the actual macrophage target for SQ109 has not been reported. These results support the proposal that macrophage stimulation contributes to the potent activity seen against intracellular *Leishmania* parasites, but not in *T. cruzi*, at least *in vitro* in Vero or LLC-MK_2_ cells. In mice models of *M. tuberculosis* infection, it was also shown that SQ109 had major effects on the immune system that contributed to its efficacy.^22^ Specifically, there were large increases in the M1-specific markers IL-6, IL-1β, TNF-α and IFN-γ and given that *T. cruzi* in animals also proliferate in macrophages, it is likely that there may be similar immune-related effects that contribute to its efficacy, in mice.^15^ It is of interest to note here that the major current therapy for leishmaniasis involves the use of miltefosine. Miltefosine has been shown to have a complex mechanism of action^34^ and in post Kala-Azar dermal leishmaniasis (PKDL)^33^ it has been shown to have both a direct antiparasitic (increased nitrite; decreased arginase activity) as well as an indirect, immunomodulatory mechanism, inducing large increases in serum IL-6 (4.7x), IL-1β (13.1x), TNF-α (4.7x) in patient macrophages and monocytes, analogous to the effects seen with SQ109 in the *M. tuberculosis*/macrophage study. Consideration of all of the results described above lead to some general proposals—a pictorial summary—for the mechanisms of action of SQ109 and its analogs, amiodarone, dronedarone, azoles, amphotericin, and miltefosine, including the roles of the host cells, as shown in Figure 7.

**Figure 7.**
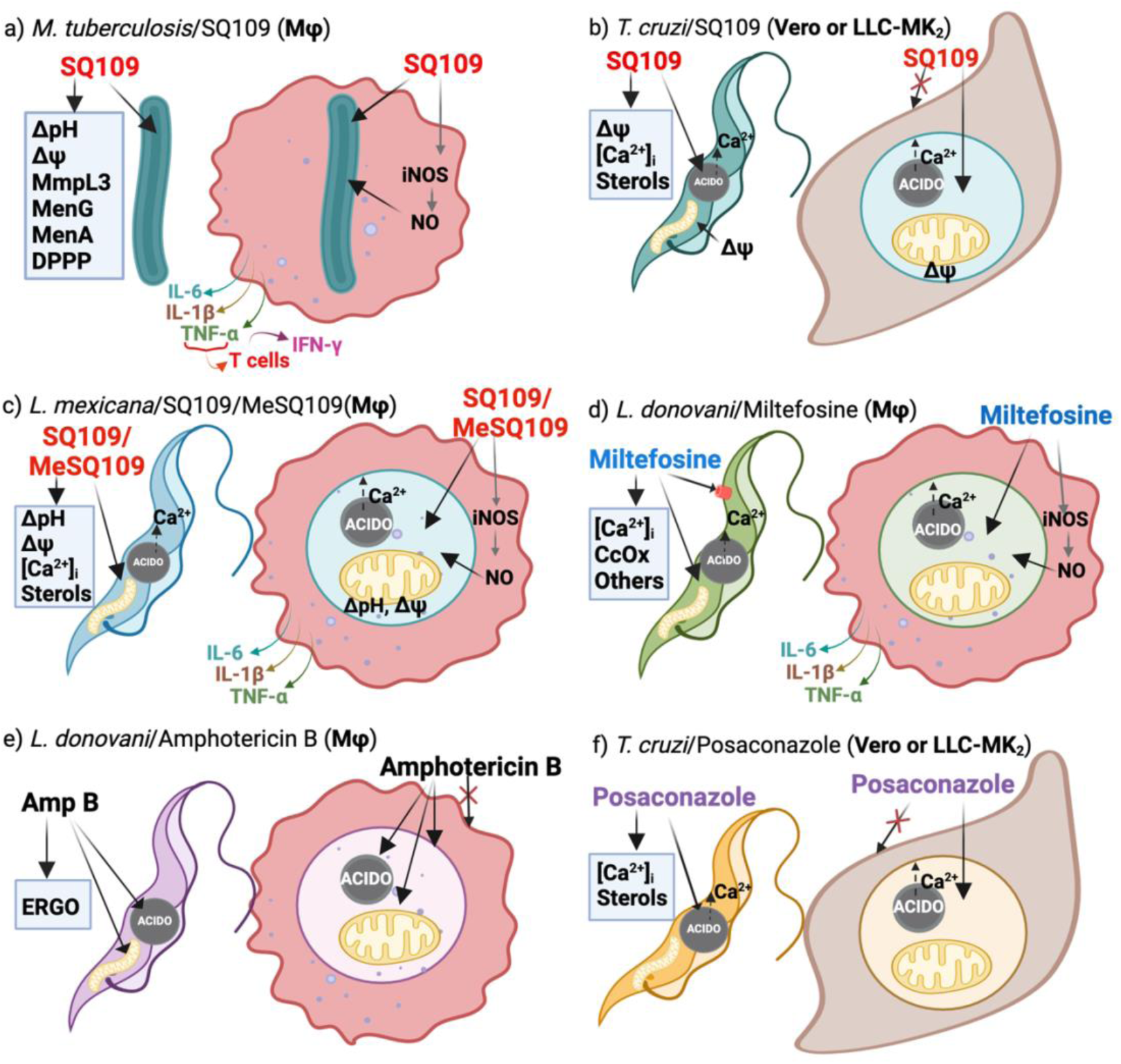
Summary of proposed mechanisms of action of SQ109, MeSQ109, azoles, miltefosine and amphotericin against *M. tuberculosis*, *Leishmania spp*. and *T. cruzi* extra- and intra-cellular forms indicating the potential role of host cells. a) *M. tuberculosis*/SQ109/macrophage (Mφ) host indicating NO/iNOS role and innate immune response. b) *T. cruzi*/SQ109/Vero (epithelial) cell host. c) *L. mexicana*/SQ109/MeSQ109/macrophage. d) *L. donovani*/miltefosine/macrophage. e) *L. donovani*/amphotericin B/macrophage. f) *T. cruzi*/posaconazole/LLC-MK_2_ (epithelial cell). Created with BioRender.com.

SQ109 itself was first developed as an agent against *M. tuberculosis* where it has been shown to target MmpL3, MenA, MenG, and undecaprenyl diphosphate phosphatase, an ortholog of the *M. tuberculosis* decaprenyl diphosphate phosphatase DPPP. It also collapses the proton motive force and there are cell growth inhibition correlations with ΔpH/Δψ in IMVs derived both from *E. coli*, as well as from *M. smegmatis*, as outlined in Figure 7a (at left). As noted above, SQ109 polarizes macrophages to the pro-inflammatory/anti-infective M1 phenotype and when inside macrophages, there are significant host cell contributions to *M. tuberculosis* cell killing involving induction of iNOS (and thus, nitric oxide). In addition, IL-1β, IL-6 and TNF-α are produced by the polarized macrophages, leading (in animals) to formation of IFN-γ, due to activation of T cells, as illustrated in Figure 7a (at right). In *T. cruzi*, Figure 7b, there are effects of SQ109 on Δψ collapse as well as on [Ca^2+^]_i_ in the extracellular epimastigotes, together with changes in sterol biosynthesis. However, for the intracellular amastigotes, *T. cruzi* is typically grown in either Vero cells or LLC-MK_2_ cells which are both epithelial cells and are not known to produce large amounts of cytokines or nitric oxide. There are differences between the IC_50_ values for SQ109, amiodarone as well as dronedarone inhibition of the extracellular promastigote form of *T. cruzi* and the intracellular form, but only about a factor of 6, Figure 7b and Table 3. In the case of *L. mexicana*, as reported previously there are collapses of the Δψ contribution to the PMF in promastigotes due to targeting of mitochondria by SQ109, together with increases in [Ca^2+^]_i_, derived from acidocalcisomes and mitochondria, Figure 7c. When grown in J774 macrophages there are much larger effects against the intracellular amastigotes, about a factor of 100, Table 3 and Figure 7c, and we propose that this has its origin, at least in part, in host cell activation, as seen with SQ109 in the *M. tuberculosis* study^22^. The downstream effects of the cytokines would, of course, be seen in animal models. With miltefosine, there is more potent activity (∼5x) against *L. donovani* promastigotes than against amastigotes, Table 3 and Figure 7d, although there is a relatively large range in the reported IC_50_ values for amastigotes, depending on the host cells used. Miltefosine has been reported to have effects on many calcium-utilizing enzymes in promastigotes^34^ as well as on other enzymes such as cytochrome c oxidase,^30^ Figure 7d, as well as on *L. donovani* infected macrophages, which express IL-1β, IL-6 and TNF-α,^23^ expected to contribute to very good *in vivo* activity. Similar effects on cytokines were observed in peripheral blood mononuclear cells in patients treated with miltefosine^23^ and it has been proposed^35^ that miltefosine-induced activation of Th1 cytokines (IFN-γ and IL-12) “is essential to prevail over the *Leishmania*-driven Th2 response”. What appears different between the effects of miltefosine and SQ109 in macrophages is that pro-inflammatory cytokines are produced in uninfected macrophages by SQ109, but not by miltefosine.^36^ In addition, miltefosine targets Akt^37^ as well as phosphatidylcholine biosynthesis.^38^

With miltefosine, the IC_50_ values against two strains of *L. donovani* were reported to be 2.2 and 5.1 μM^39^ while against amastigotes grown in macrophages there was considerable variability in activity between the host cells used: mouse peritoneal macrophages, mouse bone marrow-derived macrophages, human peripheral blood monocyte-derived macrophages (PBMΦ) and differentiated THP-1 cells.^40^ Most activity was seen with the human PBMΦs where the IC_50_ values were 0.16, 0.74 and <1.1 μM, on average <0.67 μM, while activity against amastigotes in the other three cell lines was much less, on average ∼11 μM. With amphotericin B, Figure 7e, which binds to ergosterol and disrupts cell membrane integrity, there was potent activity against amastigotes in all 4 cell lines with an average IC_50_ of 0.13 μM.^40^ Similar results for miltefosine and amphotericin were obtained by Vermeersch et al.^41^ who found only small increases in activity of amphotericin against amastigotes *versus* promastigotes, consistent with a primarily “physical” mechanism of action of amphotericin, disrupting cell membrane function, as opposed to a more host-cell-mediated effect. That notwithstanding, it is of interest to note that some of the adverse side-effects of amphotericin in infusion therapy appear to be due to cytokine release.^42^ The azole posaconazole, Figure 7f, targets ergosterol biosynthesis and hence cell membranes, as well as Ca^2+^ homeostasis, but like amphotericin, it does not act as a protonophore uncoupler: it does not collapse the pH gradient (Figure 5f) and as discussed previously, in *T. cruzi* it does not have an effect on the mitochondrial membrane potential.^4^ What seems clear is that SQ109 stimulates macrophages to release cytokines, as does miltefosine, and this is expected to contribute to intracellular cell killing via iNOS and in animals, via innate immunity: the cytokine/T-cell/INF-γ pathways. Host-mediated killing of *T. cruzi* inside Vero or LLC-MK_2_ cells would be expected to be much less, although the relatively good efficacy of SQ109 in *T. cruzi* infected mice where there is an 80% survival rate^15^, would be consistent with *T. cruzi* also growing inside macrophages in mice, leading to an increased in mice survival.

As to the effects of SQ109 on the Apicomplexan parasites, *P. falciparum* and *Toxoplasma gondii*: In *P. falciparum*, SQ109 has a ∼2-4 μM IC_50_ against the asexual blood stage growing inside red blood cells, in the range seen with *T. cruzi* epimastigotes, but there is a ∼10x increase in activity against late-stage gametocytes (which are extracellular) with an IC_50_ of 0.14 μM, an increase that may be attributed to the relatively high barrier of entry into the red cell. In *T. gondii*, the IC_50_ for the intracellular tachyzoites is 1.8 μM. These cells were grown in hTERT (human telomerase reverse transcriptase-immortalized) fibroblasts, and the modest *in vitro* activity suggests no major host-cell contribution to SQ109 activity. However, since macrophages play an important role in the early stages of *T. gondii* infection,^43^ then SQ109 may again play a role in the promising activity (80% survival) seen in a mice model of infection^15^.

## CONCLUSIONS

The results we have presented above are of interest for several reasons. First, we found that the SQ109 analog, MeSQ109, has activity (1.7 nM) against *L. mexicana* amastigotes, resident in J774 macrophages, that is a factor of ∼ 900 greater than against the extracellular, promastigote forms. There is very low activity (∼60 μM) against the macrophage host cells, resulting in a SI of ∼ 36,000. Second, we showed that activity is due at least in part to targeting of the parasite mitochondrion, collapsing the Δψ, as well as targeting acidocalcisomes, disrupting Ca^2+^ homeostasis by increasing the intracellular concentration of Ca^2+^. Third, we found that the activity of SQ109 and 17 analogs as protonophore uncouplers correlates well with cell growth inhibition in *T. brucei*, *T. cruzi* (epimastigote), *T. cruzi* (amastigote), *L. donovani*, U2OS cells as well as in *P. falciparum* with, on average, a p-value of ∼ 0.0023 for the ΔpH collapse/cell growth inhibition correlations. Fourth, we showed that the activity of SQ109 and its analogs between the different eukaryotes is highly correlated, as it is in mycobacteria, but correlations between the eukaryotes and mycobacteria are less, consistent with different additional targets. For example, MmpL3 is present in mycobacteria but is absent in eukaryotes while Ca^2+^ signaling (and sterols) are important in the eukaryotes but are absent in the bacteria. Since SQ109 has recently been shown to have a direct effect on macrophages, converting them to an M1 (pro-inflammatory) phenotype as evidenced by IL-6, IL-1β and TNF-α increases, as well as increased iNOS, this effect may contribute to the potent activity of SQ109 against *Leishmania* amastigotes, grown in macrophages, but not *T. cruzi* amastigotes, which are typically grown in monkey epithelial cells (Vero or LLC-MK_2_). However, the efficacy of SQ109 against *T. cruzi* as well as *Toxoplasma gondii* in mice models of infection^15^ may also have an immunomodulatory contribution since both cells infect macrophages, and where MAPK pathway, iNOS activation, and cytokine production may contribute to infection control, as in *M. tuberculosis* infected mice treated with SQ109. In the future, it will be of interest to test the hypothesis that M1 macrophage polarization plays a role in the potent killing of *Leishmania* spp. amastigotes, by investigating e.g. cytokine release in diverse macrophage types and with different drugs, as well as investigating the activity of MeSQ109 *in vivo*.

## METHODS

### Chemicals

SQ109, and its analogs (Figure 1) were from the batches reported previously^14^. MTT (3-[4,5-dimethylthiazol-2-yl]-2,5-diphenyltetrazolium bromide), FCCP (carbonyl cyanide-p-(trifluoromethoxy) phenylhydrazone), and nigericin were purchased from Sigma (St. Louis, MO). Acridine orange and rhodamine-123 were obtained from Molecular Probes (Eugene, OR). Dronedarone and amiodarone were from Ambeed.

### Parasites and host cell cultures

*L. mexicana* promastigotes (MHOM/BZ/1982/BEL21) were grown in liver infusion tryptose (LIT) medium, pH 7.4, supplemented with 10% fetal bovine serum and 11 mM glucose at 29 °C. J774 macrophages were grown in RPMI-1640 (Roswell Park Memorial Institute), supplemented with 10% fetal bovine serum, 2 mM L-glutamine, 100 U/mL penicillin and 100 U/mL streptomycin at 37 °C, 5% CO_2_.

### Effect of MeSQ109 on macrophages infected with *L. mexicana* amastigotes

Amastigote susceptibility assays were performed as reported (Benaim et al. 2012)^7^ with some modifications. J774 macrophages were placed over plastic coverslips inside a 24-well plate and incubated for 24 h. *L. mexicana* promastigotes were added using a ratio of 1:30 macrophages:parasites. After a 24 h incubation to ensure parasite invasion, the wells were washed twice with 1x PBS to remove any non-adherent macrophages and non-internalized parasites. Subsequently, RPMI medium with increasing concentrations of MeSQ109 was added to the wells and cells were incubated for 72 h, including appropriate controls. Coverslips were washed with PBS, fixed with methanol, and stained with Giemsa. The percentage of infected cells was determined by light microscopy.

### Determination of the mitochondrial membrane potential (Δψ_m_) of *L. mexicana* promastigotes

The fluorescent dye rhodamine-123 was used to measure the effect of SQ109 on the parasite mitochondrial membrane potential (Δψ_m_), as reported.^4^ Parasites (1.5 × 10^6^) were washed and resuspended in loading buffer (130 mM KCl, 1 mM MgCl_2_, 2 mM KH_2_PO_4_, 20 mM Tris-HCl). Subsequently, rhodamine-123 (20 μM) was added and incubated for 45 min at 29 °C under continuous stirring in the dark. Parasites were washed twice, resuspended in the same buffer, and transferred to a stirred cuvette. Measurements using different concentrations of MeSQ109 (excitation wavelength [λ_ex_] = 488 nm; emission wavelength [λ_em_] = 530 nm) were carried out using a Hitachi 7000 spectrofluorimeter at 29 °C. The protonophore FCCP (2 μM) was used as a positive control.

### Determination of the effect of MeSQ109 on acidocalcisomes from *L. mexicana* promastigotes

The effect of MeSQ109 on acidocalcisomes was performed as reported previously.^7^ Acridine orange was used as the fluorescent dye since it internalizes in acidic compartments of the cell. Promastigotes (1.5 × 10^6^) were washed and resuspended in loading buffer (130 mM KCl, 1 mM MgCl_2_, 2 mM KH_2_PO_4_, 20 mM Tris-HCl). Subsequently, acridine orange (2 μM) was added and incubated for 5 min at 29 °C under continuous stirring in the dark. Parasites were washed twice, resuspended in the same buffer, and transferred to a stirred cuvette. Measurements using different concentrations of MeSQ109 (excitation wavelength [λ_ex_], 488 nm; emission wavelength [λ_em_], 530 nm) were carried out using a Hitachi 7000 spectrofluorimeter at 29 °C. As a positive control, we used 2 μM nigericin.

### Intracellular Ca^2+^ assay in *L. mexicana* promastigotes

Intracellular Ca^2+^ was monitored as described in Rey-Cibati et al.,^8^ with some modifications. Parasites (1 × 10^8^) were loaded with Fura 2-acetoximethyl ester (Fura 2-AM) in a medium containing 145 mM NaCl, 4 mM KCl, 11 mM glucose, 2 mM CaCl_2_, 2 mM MgCl_2_, 10 mM HEPES-NaOH (pH 7.4) in the presence of Fura 2-AM (2 μM), probenecid (2.4 mM), and pluronic acid (0.05%) under continuous stirring for 2 h at 29 °C in the dark. Parasites were then washed twice with the same buffer but without Fura 2-AM. Fluorescence measurements were obtained using a Perkin-Elmer LS-55 spectrofluorimeter with a double wavelength excitation beam. Excitation was at 340 nm (Fura-2 bound to Ca^2+^) and at 380 nm (in the absence of Ca^2+^), with fluorescence emission recorded at 510 nm.

### Data analysis

GraphPad Prism 5 Software was used to calculate the half maximal inhibitory concentration (IC_50_) values. Microsoft Excel was used to calculate the mean and standard deviation.

### IMV assay

Proton translocation into IMVs was measured by the decrease of ACMA (9-amino-6-chloro-2-methoxyacridine) fluorescence, as previously reported.^17^ The excitation and emission wavelengths were 410 and 480 nm, respectively. IMVs (2 mg/mL membrane protein) were preincubated at 37 °C in 10 mM MOPS-NaOH (pH 7.0), 100 mM KCl, 5 mM MgCl_2_ containing 2 μM ACMA, and the baseline was monitored for 2 min. The reaction was then initiated by adding 1 mM ATP. When the signal had stabilized (∼6 min), compounds were added in a suitable concentration range, and proton translocation was measured fluorometrically.

## ASSOCIATED CONTENT

### Supporting Information

The Supporting Information is available free of charge at https://pubs.acs.org/doi/XXXX

Complete contact information is available at: https://pubs.acs.org/XXXX

### Author Contributions

These authors (A.M.P. and S.R.M.) contributed equally to this work. S.R.M. and A.M.P. carried out IMV assays and analyzed data. M.V-D., L. L-F., B.G.S., L.J.D-P. and A.R-C. carried out cell growth inhibition and fluorescence assays. S.H. provided IMVs and M.S. and D.S. provided inhibitors. R.B.G. and A.K. reviewed the manuscript. E.O. and G.B. wrote the paper.

### Funding Sources

This work was supported by the Fondo Nacional de Ciencia, Tecnología e Investigación, Venezuela (FONACIT Grants 2023PGP099 and 2024PGP315 to G.B.); by Chiesi Hellas (SERG grant No. 10354 to A.K.), and by a Harriet A. Harlin Professorship and the University of Illinois Foundation (E.O.)

### Notes

The authors declare no competing financial interest.

## ACKNOWLEDGEMENTS

We thank Jamie Koo, Angeline Ong, Jayannah Herdrich and Annie Le for assistance in data analysis. Some images in the TOC graphic and Figure 7 were created by the authors using BioRender.com.

## ABBREVIATIONS

U2OS: human osteosarcoma cells
EPI: *T. cruzi* epimastigotes
AMA: *T. cruzi* amastigote form
HepG2: hepatocyte carcinoma
*L. mexicana*: *Leishmania mexicana*
Mtb: *Mycobacterium tuberculosis*
MmpL3: mycobacterium membrane protein large 3
Ms: *Mycobacterium smegmatis*
MtE: *M. tuberculosis* Erdman
MtHN878: *M. tuberculosis* HN878
*P. falciparum*: *Plasmodium falciparum*
PMF: proton motive force
*T. brucei*: *Trypanosoma brucei*
*T. cruzi, Trypanosoma cruzi*: MTT, 3-(4,5-dimethylthiazol-2-yl)-2,5-diphenyltetrazolium bromide

## Notes

### Competing Interest Statement

The authors have declared no competing interest.

