## Supplementary material for "Anti-Parasitics with a Triple Threat: Targeting Parasite Enzymes, the Proton Motive Force, and Host Cell-Mediated Killing": Gustavo SI Jan 31_2025_Final.docx

**Table of Contents**

| **Table S1.** P-values for IMV IC_50_, LogP, and LogD correlations against *T. brucei*, *T. cruzi* epimastigotes (EPI), *T. cruzi* amastigotes (AMA)*, L. donovani,* and *Pf*ABS | S3 |
| --- | --- |
| **Table S2.** Correlation matrix of p-values for IMV, *T. brucei*, *P. falciparum* NF54 (ABS, asexual blood stage), U2OS, T. *cruzi* amastigotes*, T. cruzi* epimastigotes*, L. donovani* promastigotes*, M. tuberculosis* H37Rv*, M. tuberculosis* Erdman*, M. smegmatis* and *M. tuberculosis HN878* | S3 |
| **Figure S1.** Effect of MeSQ109 (**8b**) on the intracellular Ca^2+^ concentration in *L. mexicana* promastigotes. MeSQ109 at 10 μM was added (arrow) to parasites loaded with Fura-2 AM, followed by the addition of digitonin (40 μM) to reach the maximum of fluorescence, and EGTA (10 mM) was then added to obtain the minimum fluorescence, due to calcium chelation. | S4 |
| **Figure S2.** Linear correlation graphs for LogP data against *T. cruzi* EPI, *T. cruzi, L. donovani, L. mexicana, and Pf*ABS. | S5 |
| **Figure S2 (Continued).** Linear correlation graphs for LogP data against *T. cruzi* EPI, *T. cruzi, L. donovani, L. mexicana, and Pf*ABS. | S6 |
| **Figure S3.** Linear correlation graphs for LogD_7.4_ data against *T. cruzi* EPI, *T. cruzi, L. donovani, L. mexicana, and Pf*ABS. | S7 |
| **Figure S3 (Continued).** Linear correlation graphs for LogD_7.4_ data against *T. cruzi* EPI, *T. cruzi, L. donovani, L. mexicana, and Pf*ABS. | S8 |
| **Table S3.** SMILES of the compounds | S9 |

**Table S1.** P-values for IMV logIC_50_, LogP, and LogD correlations with *T. brucei*, *T. cruzi* epimastigote (EPI), *T. cruzi* amastigote (AMA)*, L. donovani* promastigote (PRO),

*L. mexicana* promastigote (PRO), and *P. falciparum* asexual blood stage (ABS) logIC_50_ values

|  | **Pearson p value** | | |
| --- | --- | --- | --- |
|  | **IMV** | **Log P** | **LogD_7.4_** |
| *T. brucei* | 0.000027 | 0.0004 | 0.629628 |
| *T. cruzi* EPI | 0.014662 | 0.111638 | 0.197737 |
| *T. cruzi* AMA | 0.008073 | 0.073407 | 0.299363 |
| *L. donovani* PRO | 0.014771 | 0.382607 | 0.138722 |
| *P. falciparum* ABS | 0.000789 | 0.001661 | 0.51239 |

**Table S2.** Correlation matrix of p-values for IMV, *T. brucei*, *P. falciparum* NF54 (ABS, asexual blood stage), U2OS, T. *cruzi* amastigotes*, T. cruzi* epimastigotes*, L. donovani* promastigotes*, M. tuberculosis* H37Rv*, M. tuberculosis* Erdman*, M. smegmatis* and *M. tuberculosis* HN878*.*

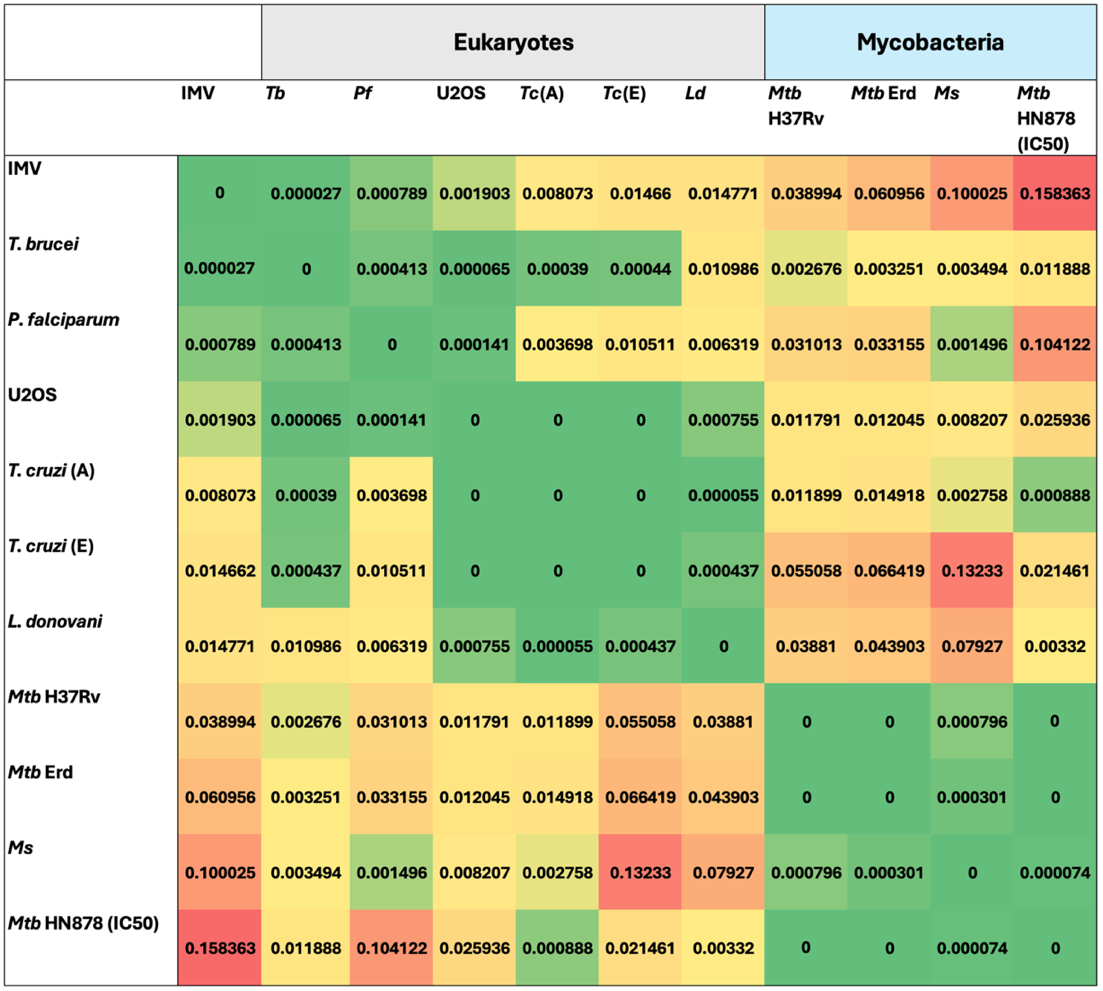

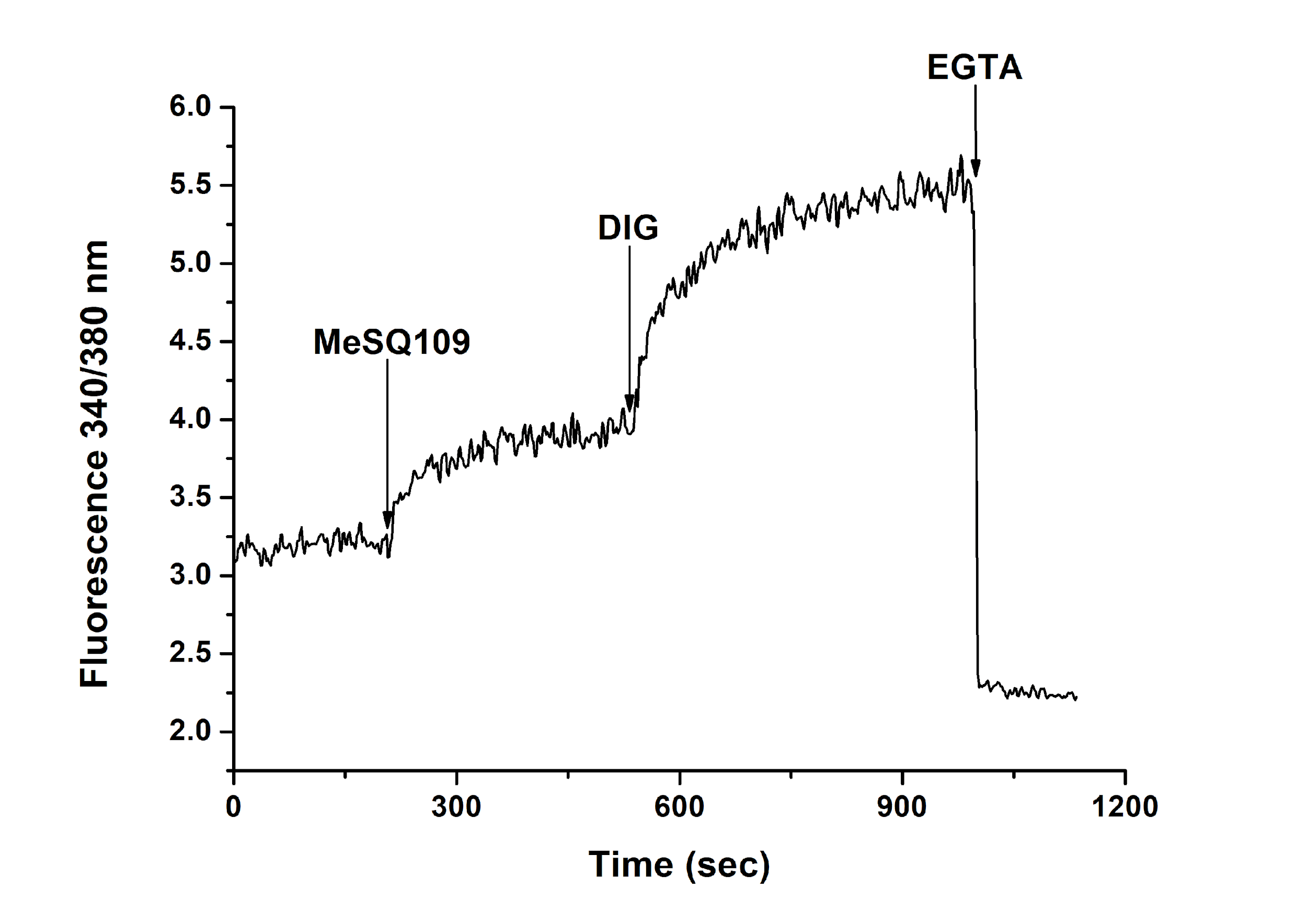

**Figure S1.** Effect of MeSQ109 (**8b**) on the intracellular Ca^2+^ concentration in *L. mexicana* promastigotes. MeSQ109 at 10 μM was added (arrow) to parasites loaded with Fura-2 AM, followed by the addition of digitonin (40 μM) to reach the maximum of fluorescence, and EGTA (10 mM) was then added to obtain the minimum fluorescence, due to calcium chelation.

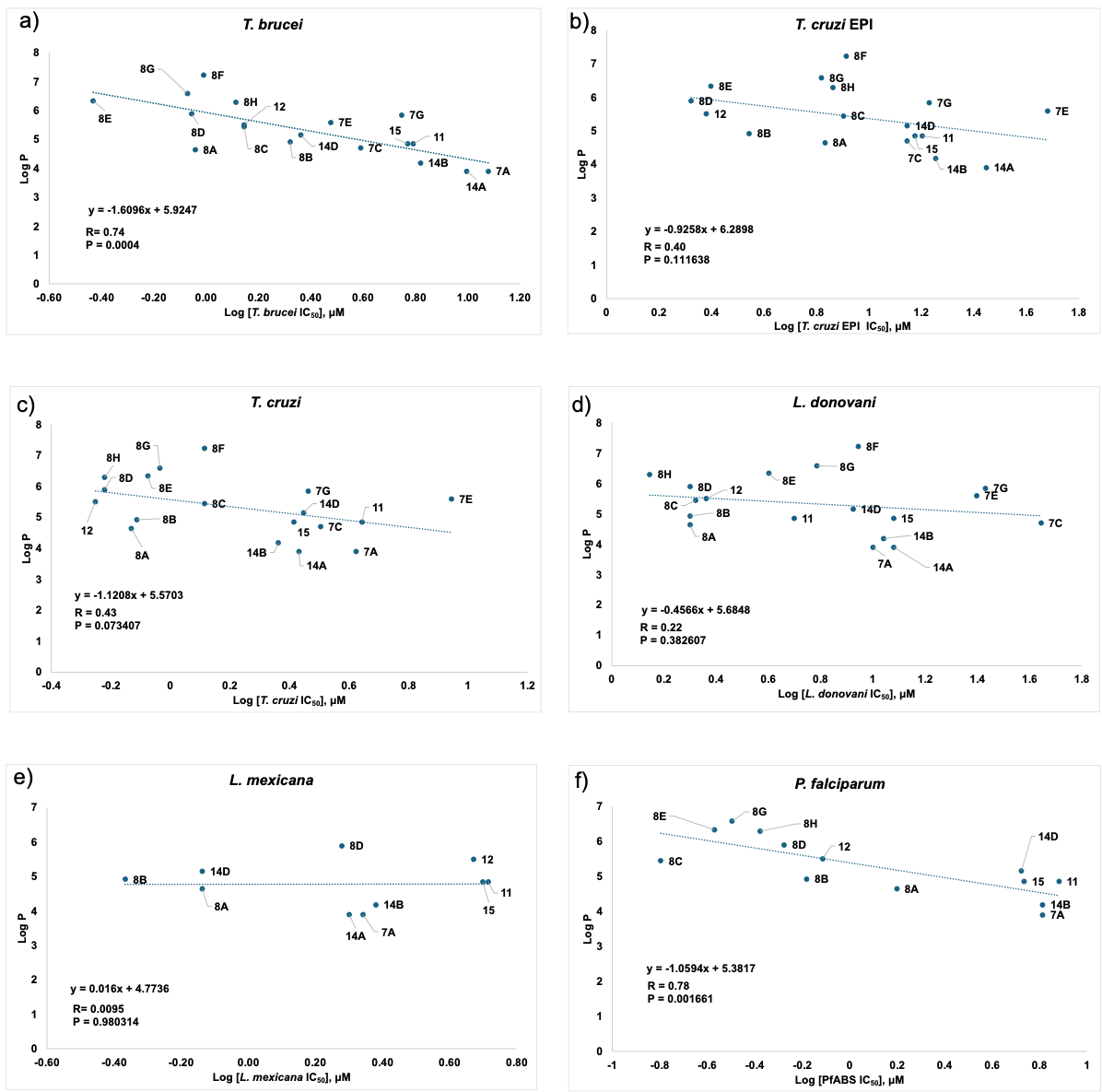

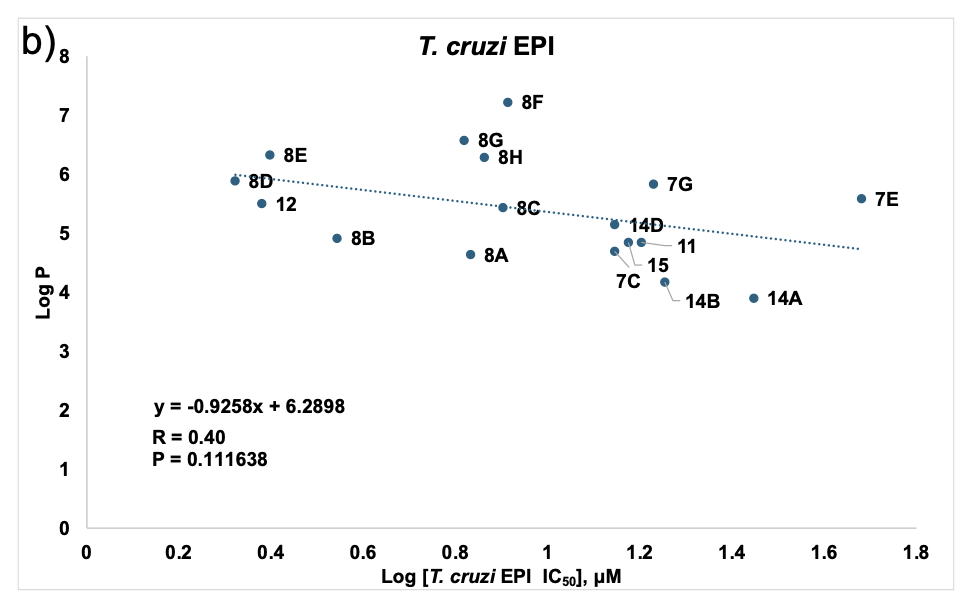

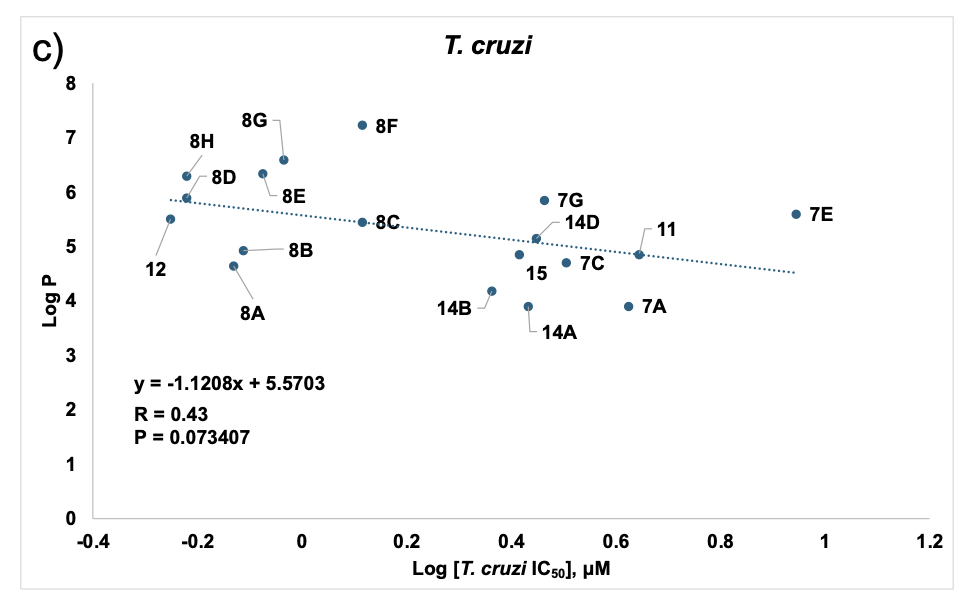

**Figure S2.** Correlations between LogP data with IC_50_ results for *T. cruzi* EPI, *T. cruzi, L. donovani, L. mexicana*, and *Pf*ABS cell growth inhibition by SQ109 and its analogs.

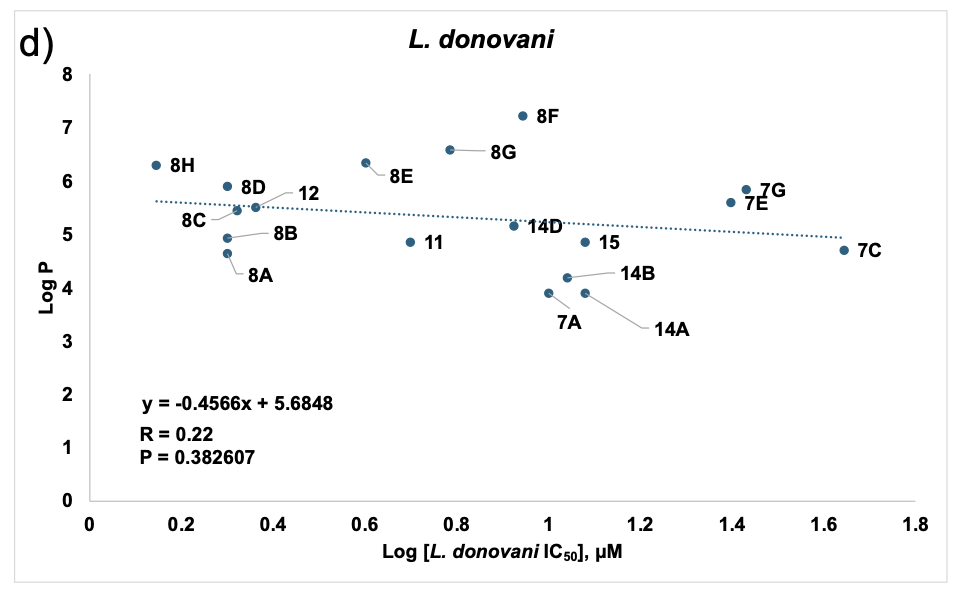

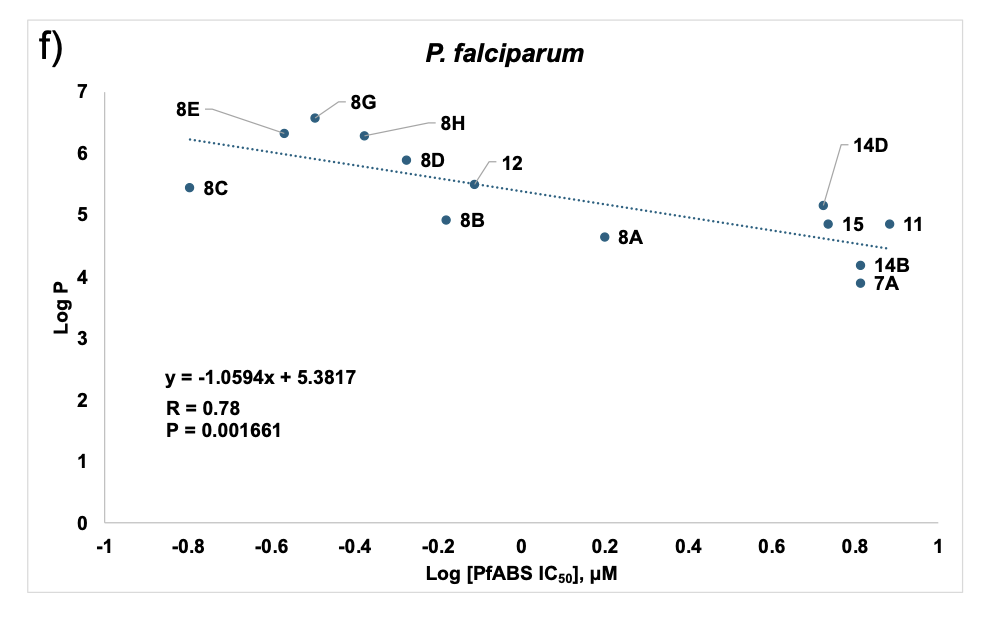

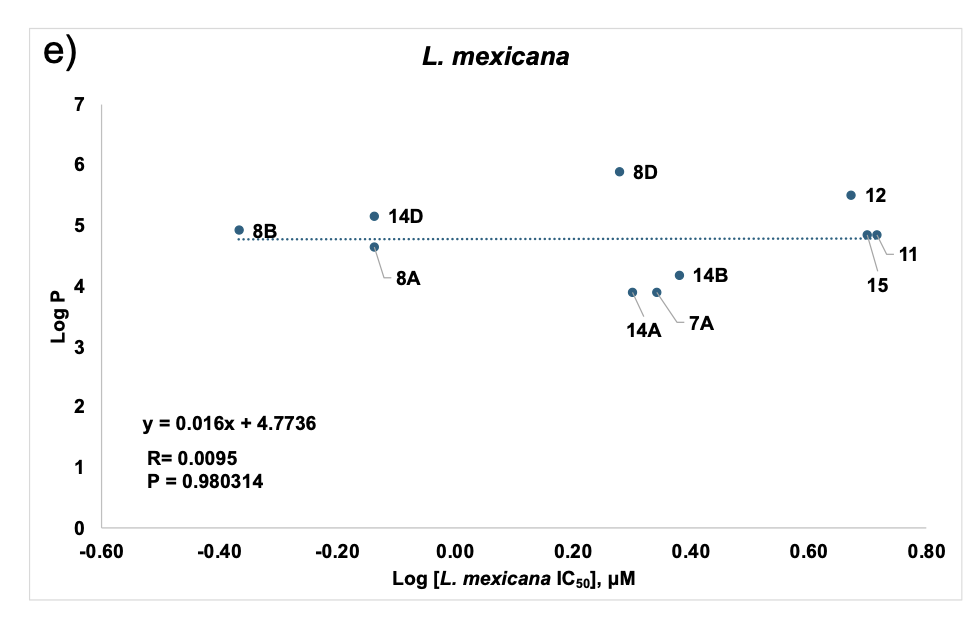

**Figure S2 (Continued).** Correlations between LogP data with IC_50_ results for *T. cruzi* EPI, *T. cruzi, L. donovani, L. mexicana*, and *Pf*ABS cell growth inhibition by SQ109 and its analogs.

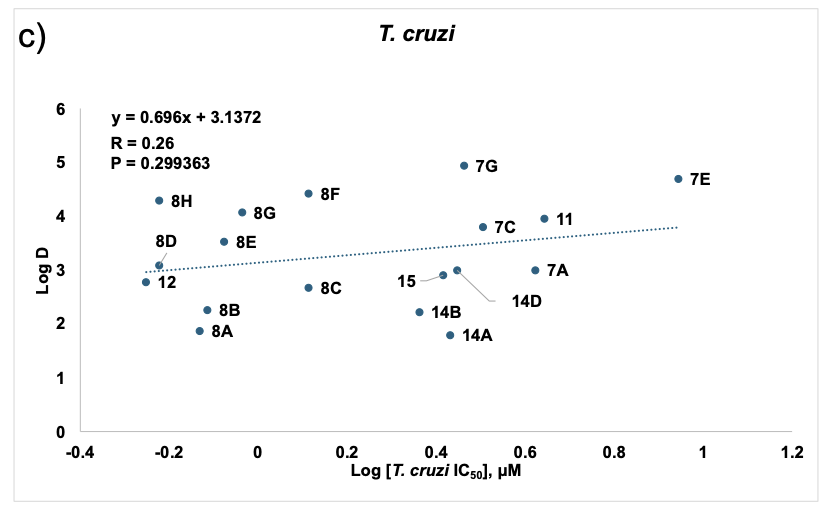

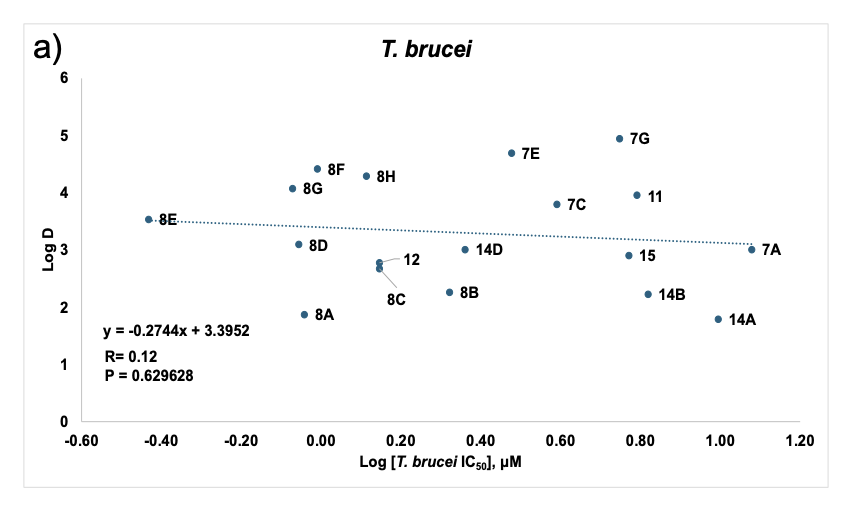

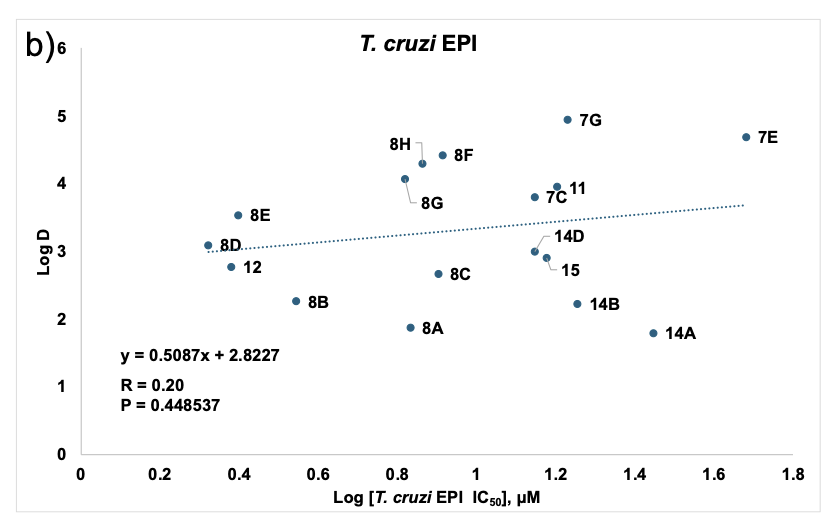

**Figure S3.** Correlations between LogD_7.4_ data with IC_50_ results for *T. cruzi* EPI, *T. cruzi, L. donovani, L. mexicana*, and *Pf*ABS cell growth inhibition for SQ109 and its analogs.

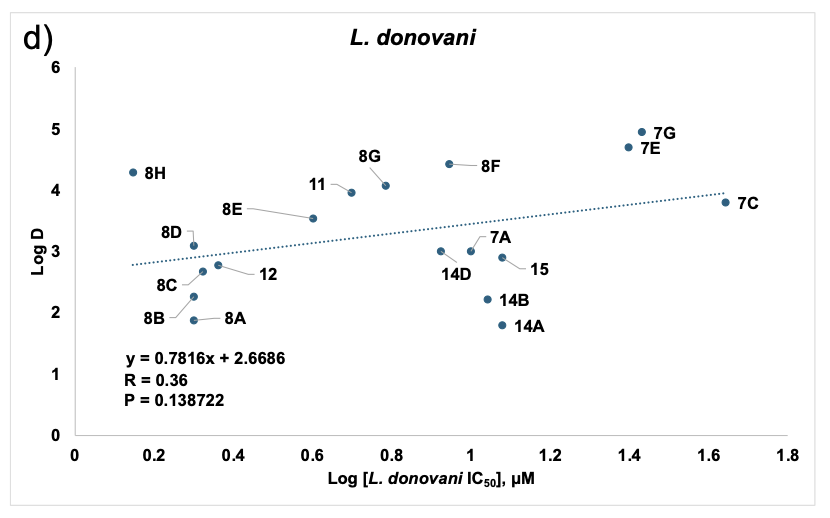

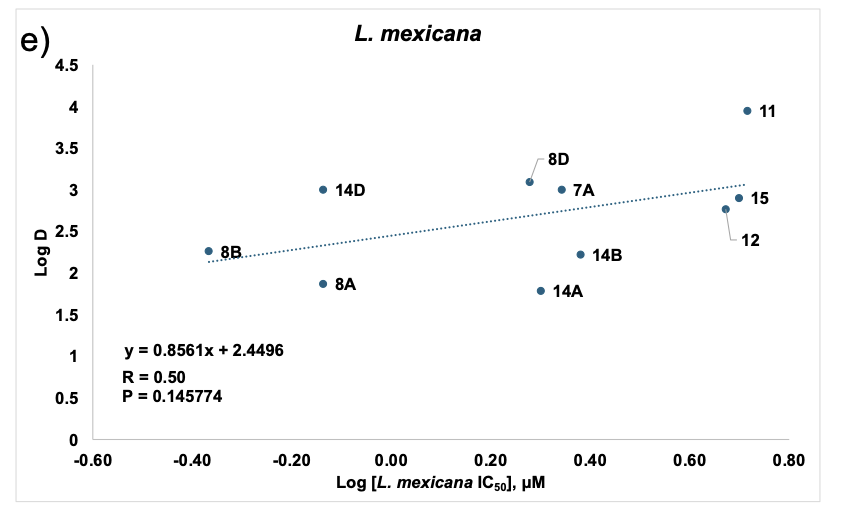

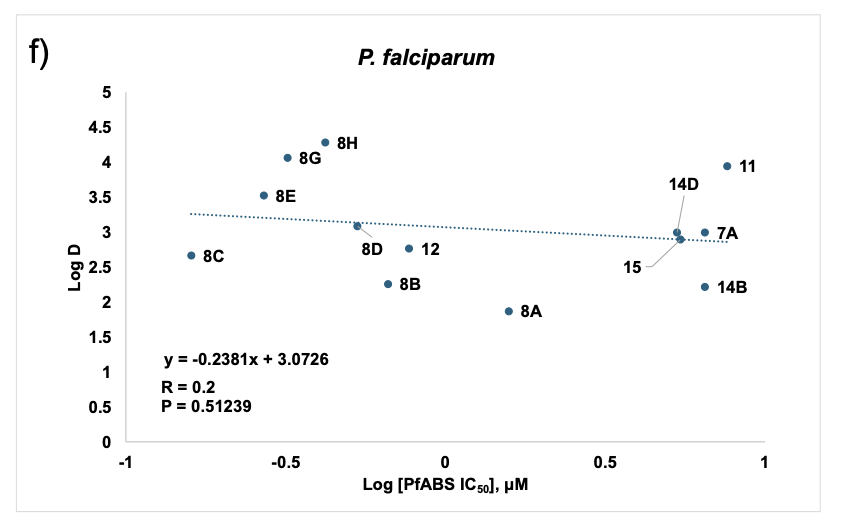

**Figure S3 (Continued).** Correlations between LogD_7.4_ data with IC_50_ results for *T. cruzi* EPI, *T. cruzi, L. donovani, L. mexicana*, and *Pf*ABS cell growth inhibition by SQ109 and its analogs.

**Table S3.** SMILES of all compounds

| **Compound** | **Structure** | **SMILES** |
| --- | --- | --- |
| **Dronedarone** | 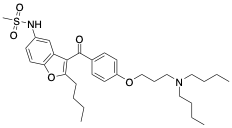 | CCCCN(CCCC)CCCOC1=CC=C(C=C1)C(=O)C1=C(CCCC)OC2=CC=C(NS(C)(=O)=O)C=C12 |
| **Amiodarone** | 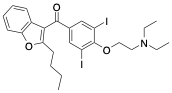 | CCCCC1=C(C(=O)C2=CC(I)=C(OCCN(CC)CC)C(I)=C2)C2=CC=CC=C2O1 |
| **8a** | 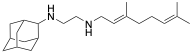 | C/C(C)=C/CC/C(C)=C/CNCCNC1[C@@H]2C[C@H](C[C@H]1C3)C[C@H]3C2 |
| **8b** | 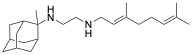 | C/C(C)=C/CC/C(C)=C/CNCCNC1(C)[C@@H]2C[C@H](C[C@H]1C3)C[C@H]3C2 |
| **8c** | 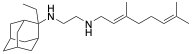 | C/C(C)=C/CC/C(C)=C/CNCCNC1(CC)[C@@H]2C[C@H](C[C@H]1C3)C[C@H]3C2 |
| **8d** | 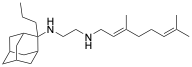 | C/C(C)=C/CC/C(C)=C/CNCCNC1(CCC)[C@@H]2C[C@H](C[C@H]1C3)C[C@H]3C2 |
| **8e** | 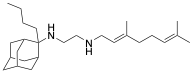 | C/C(C)=C/CC/C(C)=C/CNCCNC1(CCCC)[C@@H]2C[C@H](C[C@H]1C3)C[C@H]3C2 |
| **8f** | 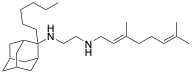 | C/C(C)=C/CC/C(C)=C/CNCCNC1(CCCCCC)[C@@H]2C[C@H](C[C@H]1C3)C[C@H]3C2 |
| **8g** | 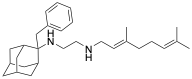 | C/C(C)=C/CC/C(C)=C/CNCCNC1(CC2=CC=CC=C2)[C@@H]3C[C@H](C[C@H]1C4)C[C@H]4C3 |
| **8h** | 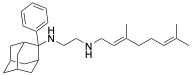 | C/C(C)=C/CC/C(C)=C/CNCCNC1(C2=CC=CC=C2)[C@@H]3C[C@H](C[C@H]1C4)C[C@H]4C3 |
| **12** | 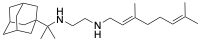 | CC(C)(C12C[C@H](C[C@@H]3C2)C[C@H](C3)C1)NCCNC/C=C(CC/C=C(C)\C)\C |
| **7a** | 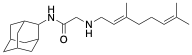 | C/C(C)=C/CC/C(C)=C/CNCC(NC1[C@@H]2C[C@H](C[C@H]1C3)C[C@H]3C2)=O |
| **7c** | 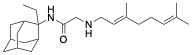 | C/C(C)=C/CC/C(C)=C/CNCC(NC1(CC)[C@@H]2C[C@H](C[C@H]1C3)C[C@H]3C2)=O |
| **7e** | 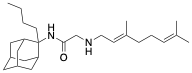 | C/C(C)=C/CC/C(C)=C/CNCC(NC1(CCCC)[C@@H]2C[C@H](C[C@H]1C3)C[C@H]3C2)=O |
| **7g** | 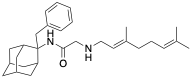 | C/C(C)=C/CC/C(C)=C/CNCC(NC1(CC2=CC=CC=C2)[C@@H]3C[C@H](C[C@H]1C4)C[C@H]4C3)=O |
| **11** | 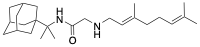 | CC(C)(C12C[C@H](C[C@@H]3C2)C[C@H](C3)C1)NC(CNC/C=C(CC/C=C(C)\C)\C)=O |
| **14a** | 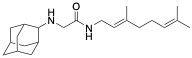 | C/C(C)=C/CC/C(C)=C/CNC(CNC1[C@@H]2C[C@H](C[C@H]1C3)C[C@H]3C2)=O |
| **14b** | 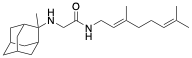 | C/C(C)=C/CC/C(C)=C/CNC(CNC1(C)[C@@H]2C[C@H](C[C@H]1C3)C[C@H]3C2)=O |
| **14d** | 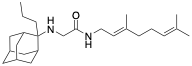 | C/C(C)=C/CC/C(C)=C/CNC(CNC1(CCC)[C@@H]2C[C@H](C[C@H]1C3)C[C@H]3C2)=O |
| **15** | 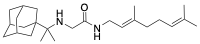 | CC(C)(C12C[C@H](C[C@@H]3C2)C[C@H](C3)C1)NCC(NC/C=C(CC/C=C(C)\C)\C)=O |
| **Miconazole** | 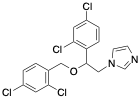 | ClC1=CC=C(COC(CN2C=CN=C2)C2=CC=C(Cl)C=C2Cl)C(Cl)=C1 |
| **Posaconazole** | 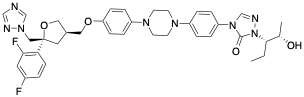 | CC[C@H](N1C(N(C2=CC=C(N3CCN(C4=CC=C(OC[C@H]5C[C@](CN6C=NC=N6)(C7=C(F)C=C(F)C=C7)OC5)C=C4)CC3)C=C2)C=N1)=O)[C@@H](O)C |
| **Amphotericin B** | 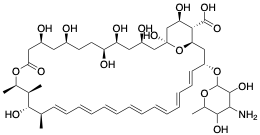 | CC1OC(O[C@@H]2C[C@@H]3O[C@@](O)(C[C@H](O)[C@H]3C(O)=O)C[C@@H](O)C[C@@H](O)[C@H](O)CC[C@@H](O)C[C@@H](O)CC(=O)O[C@@H](C)[C@H](C)[C@H](O)[C@@H](C)C=CC=C\C=C/C=C\C=CC=CC=C2)C(O)C(N)C1O |
| **Pentostam** | 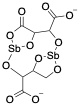 | O=C1O[Sb]2OC1C(C([O-])=O)O[Sb](O3)OCC3C(C([O-])=O)O2 |
| **Miltefosine** | 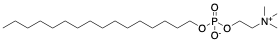 | O=C1O[Sb]2OC1C(C([O-])=O)O[Sb](O3)OCC3C(C([O-])=O)O2 |
